# Mechanism of the secretion of the lanthipeptide nisin

**DOI:** 10.1101/839423

**Authors:** Marcel Lagedroste, Jens Reiners, Sander H.J. Smits, Lutz Schmitt

**Affiliations:** Institute of Biochemistry, Heinrich Heine University Düsseldorf

**Author notes:** Center for Structural Studies, Heinrich Heine University Düsseldorf (Jens Reiners). To whom correspondence should be addressed; Lutz Schmitt, Institute of Biochemistry, Heinrich Heine University Düsseldorf, Universitätsstr. 1, 40225 Düsseldorf, Germany.

**Keywords:** Antimicrobial peptides, lanthipeptide, transport, ABC transporter, leader peptide, peptide secretion, *in vitro* activity, enzyme kinetic, RP-HPLC analysis, MS analysis

## Abstract

Lanthipeptides are ribosomally synthesized and post-translationally modified peptides containing dehydrated amino acids and (methyl-)lanthionine rings. One of the best-studied example is nisin, which is synthesized as a precursor peptide comprising of an N-terminal leader peptide and a C-terminal core peptide. Amongst others, the leader peptide is crucial for enzyme recognition and acts as a secretion signal for the ABC transporter NisT which secrets nisin in a proposed channeling mechanism. Here, we present an *in vivo* secretion analysis of this process in the presence and absence of the maturation machinery composed of the dehydratase NisB and the cyclase NisC. The data clearly demonstrated that the function of NisC, but the mere presence of NisB modulated the apparent secretion rates. Additional in vitro studies of detergent-solubilized NisT revealed how the activity of this ABC transporter is again influenced by the enzymes of the maturation machinery, but not the substrate.

## Introduction

Many natural products (NP) produced as secondary metabolites by microorganisms can be used as pharmaceuticals (e.g. as anticancer, antibacterial or antiviral drugs) (Newman and Cragg 2016). One class of these NPs are ribosomally synthesized and post-translationally modified peptides (RiPPs). The family of lanthipeptides, especially those with antimicrobial activity (lantibiotics), is gaining interest as a potential alternative for antibiotics to treat harmful multidrug resistance strains such as methicillin-resistant *Staphylococcus aureus* or vancomycin-resistant *Enterococci* (Dischinger, Basi Chipalu et al. 2014, Hudson and Mitchell 2018). Thus, it is important to gain further insights into lanthipeptide biosynthetic machineries to produce peptides with novel properties.

Lanthipeptides (LanA) are produced as precursor peptides with an N-terminal leader peptide (LP) and a C-terminal core peptide (CP) (Arnison, Bibb et al. 2013). The LP serves as a signal sequence and recognition site for the modification enzymes and the export protein (Kuipers, Beerthuyzen et al. 1993, Kuipers, de Boef et al. 2004, Mavaro, Abts et al. 2011, Abts, Montalban-Lopez et al. 2013)). Furthermore, the LP keeps the modified peptide (mLanA) inactive in the cytosol (van der Meer, Rollema et al. 1994). Additionally, the post-translational modifications (PTM) are installed within the CP and not found in the LP (van der Meer, Polman et al. 1993). These PTM’s are unusual amino acids (aa) such as didehydroalanine (Dha), didehydrobutyrine (Dhb) or (methyl-)lanthionine ((Me)Lan) (Newton, Abraham et al. 1953, Gross and Morell 1968).

Nisin is produced by the Gram-positive bacterium *Lactococcus lactis* (*L. lactis*) as a precursor peptide (pre-NisA/NisA), where the genes for modification, secretion and maturation enzymes are located on one operon (Figure 1A) (Kuipers, Beerthuyzen et al. 1993). First, the ribosomal synthesized NisA is modified by the modification enzymes NisB and NisC (Figure 1B, I). Within the unmodified pre-NisA (uNisA) serine and threonine residues are dehydrated by the dehydratase NisB via a tRNA-depended glutamylation and elimination reaction to Dha and Dhb residues (Karakas Sen, Narbad et al. 1999, Garg, Salazar-Ocampo et al. 2013, Ortega, Hao et al. 2015). Subsequently, the dehydrated aa are coupled to neighboring cysteine residues via a Michael-like addition catalyzed stereo- and regio-specifically by the cyclase NisC (Koponen, Tolonen et al. 2002, Okeley, Paul et al. 2003, Li and van der Donk 2007). The reaction of both enzymes follows an alternating mode with a N- to C-terminus directionality yielding (Me)Lan residues (Lubelski, Khusainov et al. 2009, Repka, Hetrick et al. 2018). Next, the exporter protein NisT secrets the modified NisA (mNisA) to the exterior (Figure 1B, II) (Quiao and Saris 1996). Finally, the LP is cleaved by the extracellular located serine protease NisP and active nisin is released (Figure 1B, III) (van der Meer, Polman et al. 1993).

**Figure 1:**
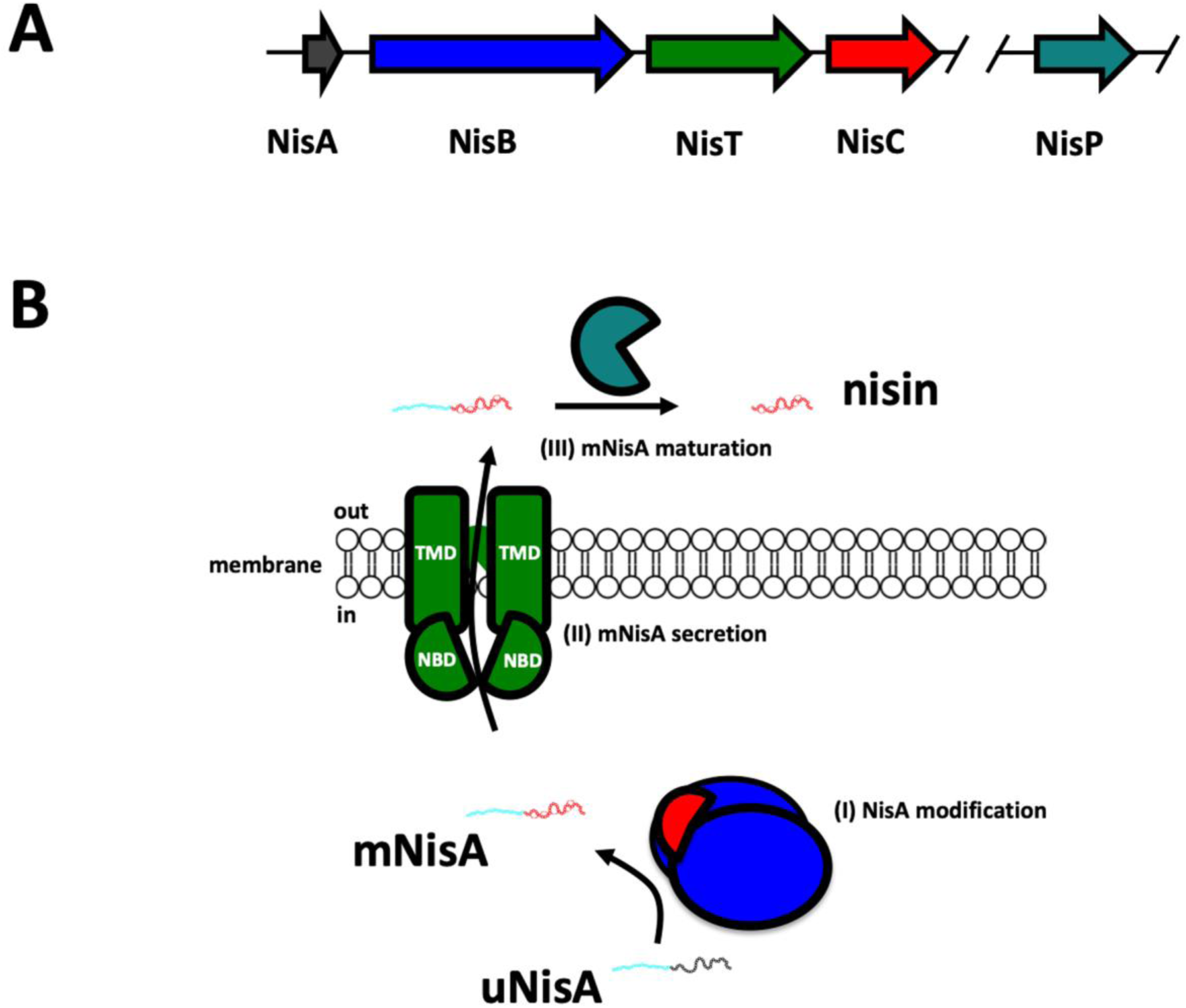
Scheme of nisin modification and secretion system. (**A**) The lanthipeptide nisin (NisA, grey) operon encodes for the modification and secretion enzymes. (**B**) The enzymes NisB (blue) catalyze the dehydration reaction of unmodified NisA (uNisA), whereas NisC (red) catalyze the thioether ring formation resulting in modified NisA (mNisA). The ABC transporter NisT (green) translocates mNisA across the membrane to the exterior. Finally, the mature peptide is processed by the serine protease NisP (light grey) and active nisin is released. Please note, that the operon is partial represented and shows only proteins responsible for nisin maturation and secretion.

The lanthipeptide exporters (LanT) belong to the superfamily of ABC transporters, which are found in all kingdoms of life (Higgins 1992). Bacterial ABC transporters comprise mainly of two domains (Fath and Kolter 1993). One domain is the transmembrane domain (TMD) creating the translocation tunnel. The other domain is the nucleotide-binding domain (NBD), which binds and hydrolyses ATP to energizes conformational changes used for substrate translocation. Some lanthipeptide exporters harbor an additional domain, a C39 peptidase domain. This domain is classified as a bacteriocin-processing endopeptidase (Havarstein, Diep et al. 1995) and we term this subfamily of transporters LanT_C39P_.

All known lanthipeptide exporters function as dimers and translocation of the lanthipeptide is LP-dependent (Klein, Kaletta et al. 1992, Schnell, Engelke et al. 1992, Bierbaum, Brotz et al. 1995, Kuipers, de Boef et al. 2004). For some LanT/ Lan_C39P_T proteins it is proposed that the exporter and modification proteins assemble a multimeric enzyme complex at the membrane and is translocated to the exterior (e.g. nisin, subtilin and nukacin ISK-1) (Siegers, Heinzmann et al. 1996, Kiesau, Eikmanns et al. 1997, Nagao, Aso et al. 2005).

In 2004, it was shown that the secretion of nisin without the modification enzymes NisB, NisC and the protease NisP was possible (Kuipers, de Boef et al. 2004). *In vivo* studies on the secretion process expanded the knowledge on the nisin modification and secretion system (Kluskens, Kuipers et al. 2005, Rink, Kuipers et al. 2005, Rink, Kluskens et al. 2007, Kuipers, Meijer-Wierenga et al. 2008). According to these studies the observed high secretion efficiency of NisA by NisBTC was explained by a ‘channeling mechanism’ (van den Berg van Saparoea, Bakkes et al. 2008, Lubelski, Khusainov et al. 2009). Other studies focused on the application of the nisin modification machinery to produce nisin variants or lantibiotics and secrete them by NisT (Zhou, van Heel et al. 2015, van Heel, Kloosterman et al. 2016, Zhou, van Heel et al. 2016, Lagedroste, Reiners et al. 2019). Despite the *in vivo* analysis of pre-NisA secretion, only a qualitative analysis of the supernatants was performed (van den Berg van Saparoea, Bakkes et al. 2008). Thus, a systematic and quantitative analysis of the secretion mechanism by determining kinetic parameter for pre-NisA translocation by NisT is still required.

In our study, we performed an *in vivo* and *in vitro* characterization of NisT to shed light on the secretion mechanism of pre-NisA. We determined the kinetic parameter for the pre-NisA secretion by analyzing the supernatant of pre-NisA secreting *L. lactis* strains via RP-HPLC. The resulting apparent secretion rate (NisA•NisT^−1^•min^−1^) of NisT was compared with the rate of the NisBTC system and demonstrated a large enhancement in the presence of the modification machinery. The *in vitro* characterization of NisT is the first study revealing insights into the specific activity of a LanT lanthipeptide transporter and its modification enzymes as well as its substrate. In conclusion, we demonstrate an enhancement of the secretion through the maturation enzymes and a pivotal and bridging function of the dehydratase NisB in the interaction of NisC and NisT.

## Results

### *In vivo* secretion assay of pre-NisA

To obtain further insights into the mechanism of lanthipeptide secretion, the pre-NisA secretion level of the *L. lactis* strain NZ9000 was investigated in the presence and absence of the modification machinery. Here, the well-known nisin secretion and maturation system (Rink, Kuipers et al. 2005, van den Berg van Saparoea, Bakkes et al. 2008, Lubelski, Khusainov et al. 2009, van Heel, Kloosterman et al. 2016) was used to establish an *in vivo* secretion assay, where the supernatants were employed to determine the secretion level of pre-NisA peptide via RP-HPLC analysis. In our study, we used the strain *L. lactis* NZ9000BTC (Supplementary File 1) producing fully modified pre-NisA (mNisA) that is secreted by NisT. The secretion of other modification states of pre-NisA can be analyzed by deletions of one of the modification enzymes or by creating inactive mutants. The mutation NisC_H331A_ (strain NZ9000BTC_H331A_) or the deletion of NisC (strain NZ9000BT) resulted in the secretion of dehydrated NisA (dNisA). Deletion of NisB (strain NZ9000TC) or NisB and NisC (strain NZ9000T) resulted in the secretion of unmodified NisA (uNisA). The deletion of NisT (strain NZ9000BC) or the mutation of the histidine of the H-loop in the NBD of NisT (strain NZ9000BT_H551A_C) totally abolished pre-NisA secretion (Figure 2). The latter two strains were defined as the background of the secretion analysis, in which pre-NisA is expressed but not secreted. For the secretion analysis, all *L. lactis* strains were grown in minimal medium at 30°C and samples of supernatants were analyzed after induction every hour (0-6 h). Subsequently, the supernatant was analyzed by RP-HPLC (Figure 2, Supplementary Figure 1) and the peak area of pre-NisA peptides was determined to calculate the amount of peptide. The amount of secreted peptide was plotted against time and a non-linear fitting was applied to determine V_max_ (maximal amount of peptide; in nmol) and K_0.5_ (time point of 50% secreted peptide in min). These kinetic parameters allowed a direct comparison of the secretion efficiency of the different *L. lactis* strains employed in this study.

**Figure 2:**
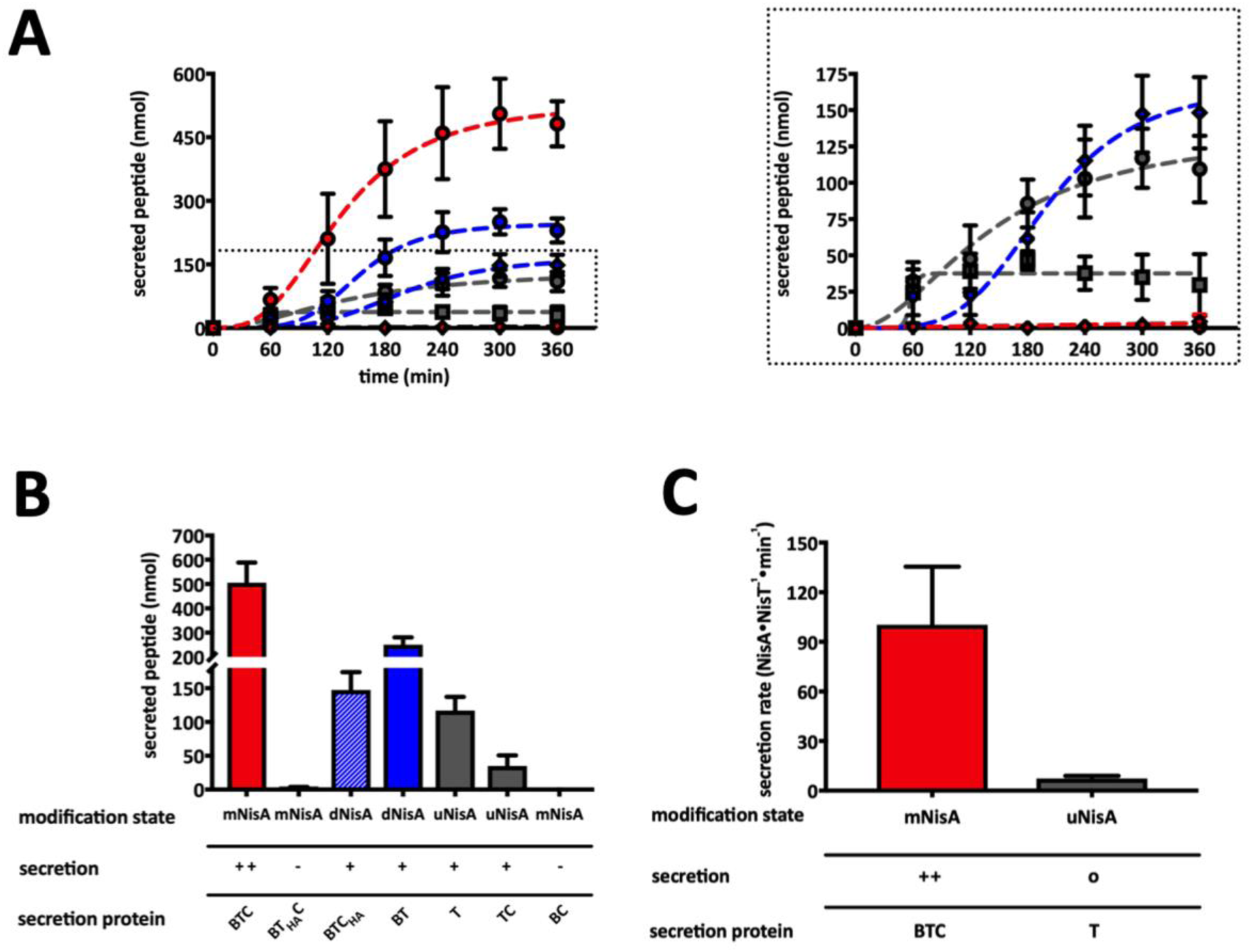
*In vivo* secretion assay of different *L. lactis* NZ9000 strains. (**A**) The supernatants of pre-NisA secreting *L. lactis* NZ9000 strains was analyzed by RP-HPLC and the amount of pre-NisA was determined. Amounts of secreted pre-peptides (nmol) are plotted against time (min) and the resulting curves were fitted by an allosteric sigmoidal fit. Modified NisA (mNisA, red) was secreted by strain NZ9000BTC (red dots) and can be preclude by *nisT* deletion (strain NZ9000BC, clear dot) or an ATP-deficient mutant (NZ9000BT_H551A_C, red rhomb). Dehydrated NisA (dNisA, blue) was secreted by strains NZ9000BTC_H331A_ (blue rhomb) and NZ9000BT (blue dot), whereas unmodified NisA (uNisA, grey) was secreted by the strains NZ9000T (grey dots) and NZ9000TC (grey square). Dashed square shows a zoom-in on strains with lower secretion level. (**B**) The kinetic parameter of V_max_ (nmol) of secreted peptides was plotted as bars against the various secretion systems. (**C**) The secretion rate of NisA molecules per NisT molecules was plotted against time (min) and fitted by linear regression. The slope represented the secretion rate of NisA•NisT^−1^•min^−1^ for the strains NZ9000BTC and NZ9000T. All data represent secretion experiments from at least five different transformants and are represented as means ± SD (n=5). ++: WT secretion; o: low secretion; −: no secretion

Strain NZ9000BTC had a V_max_ value of 534 ± 44 nmol and a K_0.5_ of 134 ± 12 min. It secreted pre-NisA most efficiently in comparison to the other strains (Figure 2, Supplementary File 3). In the cytoplasmic fraction of this strain only a small amount of mNisA was detected by WB for the first three hours after induction (Supplementary Figure 2A). This finding is similar to the previously published data for the nisin secretion and modification system (van den Berg van Saparoea, Bakkes et al. 2008).

Mutation of the H-loop at position 551 to alanine in the NBD of the ABC transporter NisT abolished the secretion of mNisA and no peptide was detected in the supernatant (Figure 2, Supplementary Figure 2C). The same result was observed for the strain NZ9000BC (Supplementary Figure 2G). Here, high amounts of mNisA were detected in the cytoplasmic fraction of the strains NZ9000BT_H551A_C and NZ9000BC (Supplementary Figure 1C/G). Thus, *nisT* deletion and the H-loop mutation both abolished mNisA secretion.

Deletion of *nisC* (strain NZ9000BT) resulted in a lower secretion efficiency of pre-NisA and the V_max_ value of 247 ± 15 nmol and a K_0.5_ of 152 ± 9 min (Figure 2, Supplementary File 3). The amount of secreted dNisA was reduced by a factor of 2.2 in comparison to mNisA (strain NZ9000BTC). Interestingly, the mutation of the catalytic histidine residue (H331) to alanine (Li, Yu et al. 2006) reduced the V_max_ value (168 ± 16 nmol) by an additional factor of 3.2. The K_0.5_ value increased to 200 ± 16 min (Figure 2). The analysis of the cytoplasmic fraction of the strain NZ9000BTC_H331A_ showed a higher amount of pre-NisA inside the cell, which only slowly decreased over the time (Supplementary Figure 2B). Slightly lower amounts of pre-NisA were observed in the cytoplasmic fraction of NZ9000BT (Supplementary Figure 2D).

Strain NZ9000T, which was obtained after the deletion of *nisB* and *nisC*, had a slightly reduced V_max_ value of 137 ± 30 nmol with a K_0.5_ of 144 ± 41 min (Figure 2, Supplementary File 3). The secretion of pre-NisA was reduced by a factor of 3.9 compared to strain NZ9000BTC. The lowest amount of secreted peptide was determined in the supernatant of strain NZ9000TC with a V_max_ value of 38 ± 8 nmol. Here, a higher amount of uNisA was detected in the cytoplasmic fraction, whereas no peptide was observed in strain NZ9000T.

In all strains, NisB, NisC and NisT were detected in their corresponding fraction (cytoplasmic or membrane) (Supplementary Figure 2A-G). The proteins NisB, NisC/NisC_H331A_ and pre-NisA were observed in the cytoplasmic fractions (except uNisA from NZ9000T). NisT was detected by WB in the membrane fraction of all strains (Supplementary Figure 2A-G). In the case of mNisA expressing strains (NZ9000BTC, NZ9000BT_H551A_C, and NZ9000BC) all proteins were detected by WB even at time point zero.

### Determination of the apparent secretion rate of mNisA

We measured the apparent secretion rate (V_S app._) by an *in vivo* secretion assay. First, the amount of secreted mNisA at different time points was determined and plotted as nmol mNisA against time (Figure 2). Second, the amount of the ABC transporter NisT was obtained by analyzing the membrane fraction of strains NZ9000BTC and NZ9000T at each time point by WB (Supplementary Figure 3A-B). Here, known concentrations of purified NBD protein were used as a standard to determine the amount of NisT on pmol (Supplementary Figure 3C). Combing these data, the secretion rate of NisA molecules per NisT molecules was calculated. The plot of nmol NisA•NisT^−1^ against time (min) was fitted by linear regression (Supplementary Figure 3D). The slope of the linear regression corresponds to V_S app._ of NisA•NisT^−1^•min^−1^, which is 100.3 ± 35.2 for the strain NZ9000BTC (Figure 2C). A strongly reduced secretion rate was determined for the strain NZ9000T (factor of 7.4 ± 1.6) as described qualitatively previously (van den Berg van Saparoea, Bakkes et al. 2008).

However, the functional unit of NisT is a dimer. Thus, the rate is twice as high as calculated (200.6 ± 70.4 NisA•NisT ^−1^•min^−1^). If one now considers that one mNisA molecule consists of 57 amino acids, the V_S app._ can be further expressed as amino acids (aa) per NisT dimer per second. Here, the V_S app._ value of the nisin exporter NisT in strain NZ9000BTC is 190.6 ± 66.8 aa•s^−1^, whereas the secretion rate of strain NZ9000T was reduced to approximately 7% (13.9 ± 3.1 aa•s^−1^). The determined secretion rate thus clearly demonstrated an enhancement in the presence of the modification enzymes NisB and NisC.

### Purification and basal ATPase activity of NisT, a class I lanthipeptide transporter

In order to determine the *in vitro* activity of a lanthipeptide ABC transporters, we purified NisT the exporter of the class I lanthipeptide nisin as a deca-histidine tagged protein variant (10HNisT). 10HNisT was homologously expressed in *L. lactis* NZ9000 and purified to high purity (≥95% as estimated by Coomassie brilliant blue stained SDS-PAGE, Figure 3A) after solubilization with the lipid-like surfactant Fos-choline-16 (FC-16, Anatrace) by immobilized metal ion affinity (IMAC) and size-exclusion chromatography (SEC). Similar to other membrane proteins, 10HNisT (72.5 kDa) showed a higher mobility on the SDS-PAGE gel and migrated at approximately 60 kDa. 10HNisT eluted as a homogeneous peak from SEC (Figure 3A), subsequently the main elution fractions were further concentrated to 50 µM and used for ATPase activity assay.

**Figure 3:**
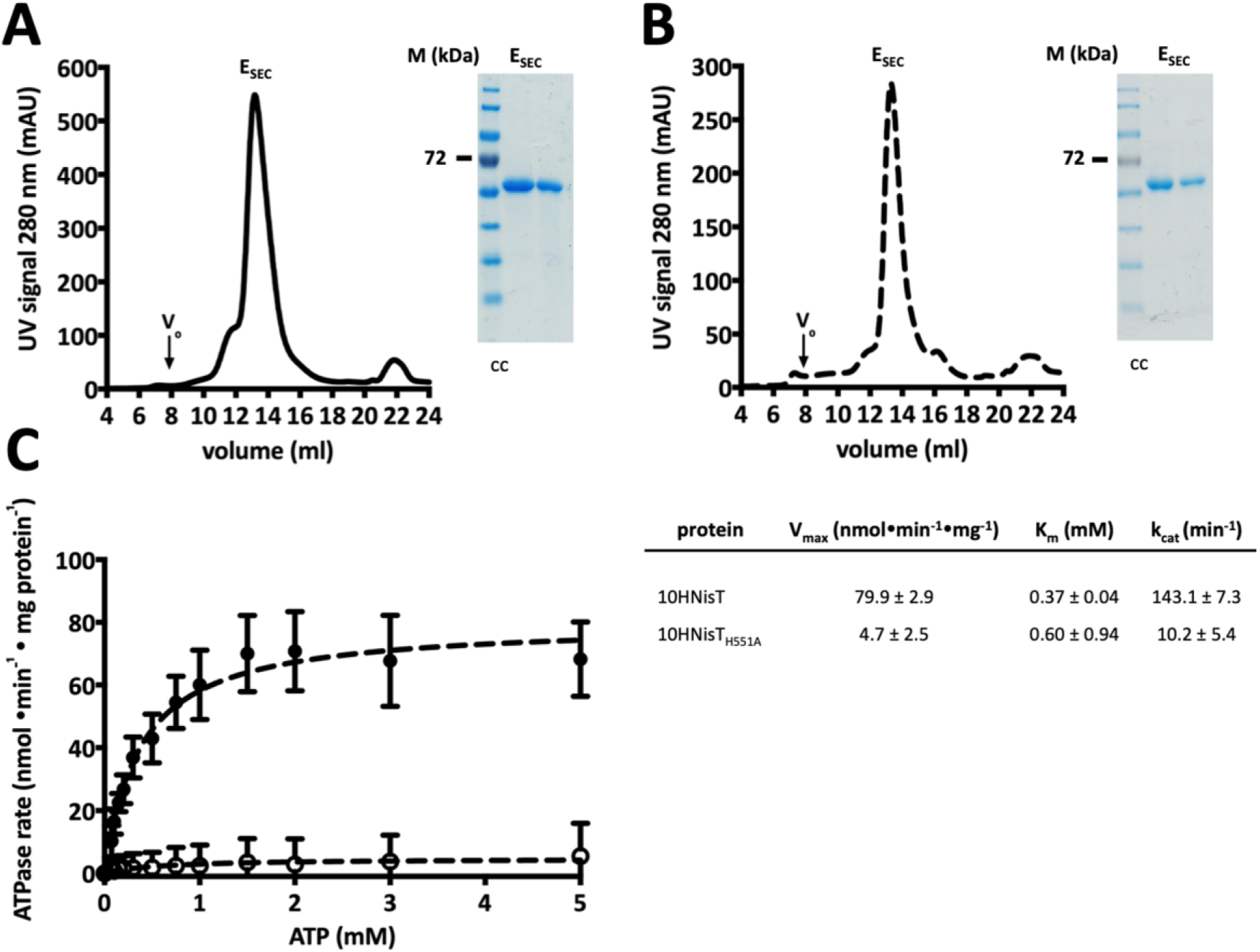
Purification and ATPase activity assay of NisT. (**A**) SEC chromatogram of 10HNisT (WT, black line) displayed a homogeneous peak (E_SEC_) at 13 ml on a Superose 6 10/300 GL column (V_0_: void volume of the column). Inset: A typical colloidal Coomassie (cc) stained SDS-PAGE gel shows a protein band between 55 and 72 kDa marker protein bands (M). (**B**) Purification of the H-loop mutant 10HNisT_H551A_ (HA, dashed line) showed comparable results for SEC profile and SDS-PAGE gel (inset). (**C**) The specific ATPase rate (nmol•min^−1^•mg protein^−1^) of purified WT (black dot) and HA mutant (unfilled circle) was plotted against ATP concentration (mM) to determine kinetic parameters. The ATPase rate was fitted by Michaelis–Menten equation to determine V_max_ (nmol•min^−1^•mg protein^−1^), K_m_ (mM) and k_cat_ (min^−1^). Activity assays were performed from five independent experiments with three replicates and are represented as means ± SD (n=5).

For the ATPase activity assay, the detergent was exchanged to CYMAL5 (Anatrace) and the ATPase rate was expressed as specific ATPase rate (nmol•min^−1^•mg^−1^). The kinetic parameters of 10HNisT in detergent solution were determined and resulted in V_max_, K_m_ and k_cat_ values for the transporter without its substrate (basal ATPase activity). The concentration of 10HNisT was kept constant (1 µM), whereas the concentration of ATP was varied from 0 to 5 mM and the reaction was stopped after 30 min. The basal ATPase rate of 10HTNisT had a V_max_ value of 79.9 ± 2.9 nmol•min^−1^•mg^−1^, a K_m_ value of 0.37 ± 0.04 mM resulting in a k_cat_ value of 143.1 ± 7.3 min^−1^ (Figure 3C). As a control the H-loop mutant of 10HNisT (10HNisT_H551A_; HA-mutant) was also purified following the same protocol and used in the ATPase activity assay (Figure 3B). The ATPase rate of the HA-mutant was reduced by a factor of 17 (V_max_ value 4.7 ± 2.5 nmol•min^−1^•mg^−1^). The K_m_ value increased by a factor 1.62 (0.60 ± 0.94 mM), whereas the k_cat_ value was 10.2 ± 5.4 min^−1^ (14-fold lower than WT 10HNisT) (Figure 3C).

### *In vitro* ATPase activity with pre-NisA variants

To investigate the effect of substrate on the ATPase rate of 10HNisT, we added different pre-NisA variants. First, the pre-NisA peptides in different modification states (uNisA, dNisA and mNisA, respectively) were purified (Supplementary Figure 4). Additionally, the leader peptide of NisA (NisA_LP_) was used in the ATPase assay to evaluate whether the isolated LP is sufficient for recognition by NisT.

For the activity assay the ATP concentration was kept constant at 5 mM, while the substrate concentration was varied from 0 to 40 µM. 10HNisT was pre-incubated with the peptides prior to the activity assay. The basal activity of 10HNisT was set to 100% and the ATPase rate with substrates was expressed as normalized ATPase rate. The ATPase rate of 10HNisT was slightly increased for all peptides. Some values like 20 µM uNisA showed a stronger stimulation of the transporter (140 %, Figure 4B), while other (e.g. 20 µM mNisA) showed a lower stimulating effect (110%, Figure 4D). However, a concentration dependent stimulation of the transporter was not observed for all the different tested peptides (Figure 4A-D) suggesting that the ATPase is not modulated by the pre-NisA variants.

**Figure 4:**
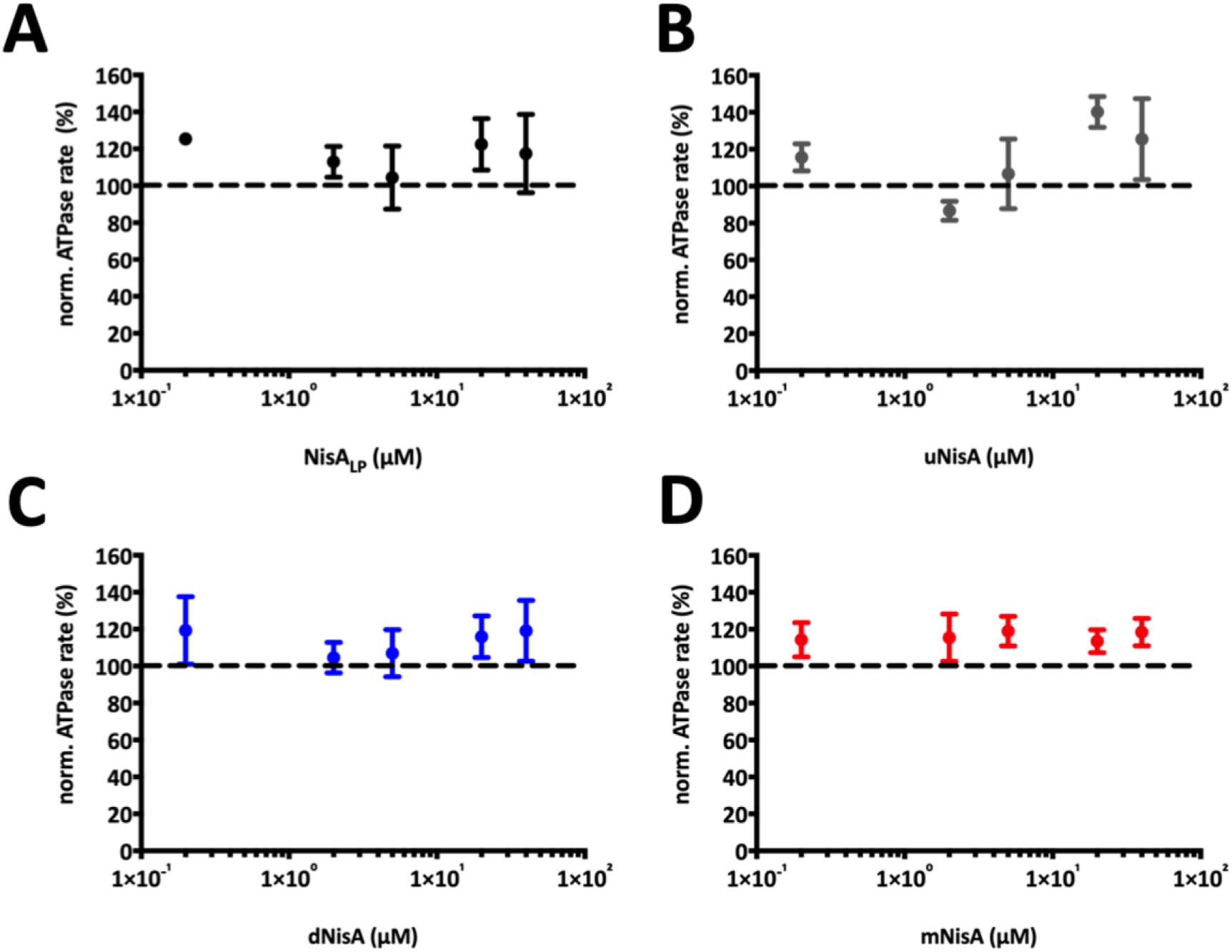
Dependence of NisT ATPase rate on different substrates. The ATPase rate of purified 10HNisT was analyzed in the presence of different substrates. (**A**) The leader peptide of NisA (NisA_LP_, black dots), (**B**) uNisA (grey dots), (**C**) dNisA (blue dots) and (**D**) mNisA (red dots) was used in various concentrations (µM) and the ATPase rate is shown as normalized ATPase rate (%). The basal ATPase rate was set as 100% (dashed line) and further values were normalized accordingly. The assays were performed in at least four independent experiments and are represented as means ± SD (n=4).

### *In vitro* ATPase activity in presence of NisBC and mNisA

Previous studies (van den Berg van Saparoea, Bakkes et al. 2008) and our data demonstrated that the secretion of pre-NisA is strongly enhanced by the modification proteins NisB and NisC (see *in vivo* secretion assay) and therefore the ATPase rate of NisT might also be influenced by these interaction partner. To investigate the effect of NisB and NisC on the ATPase rate of 10HNisT the ATPase activity assay was repeated under the same conditions (ATP concentration constant, various concentration of interaction partner). We observed that the ATPase rate of NisT was independent of the various concentrations of NisB or NisC. Thus, only fixed molar ratio of 10HNisT to the interaction partner was used (NisT:NisB/NisC 1:2; in the case of NisT:NisBC 1:2:2) (Figure 5A). The basal ATPase rate of NisT was 62.5 ± 9.4 nmol•min^−1^•mg^−1^ and was not changed within the experimental error in the presence of NisB or NisC (54.7 ± 4.5 nmol•min^−1^•mg^−1^, 59.3 ± 4.9 nmol•min^−1^•mg^−1^). If both proteins were used in the assay (NisBC), the ATPase rate of 10HNisT was reduced by a factor of 1.3 (49.3 ± 4.4 nmol•min^−1^•mg^−1^), but the difference was not significant (Figure 5A).

**Figure 5:**
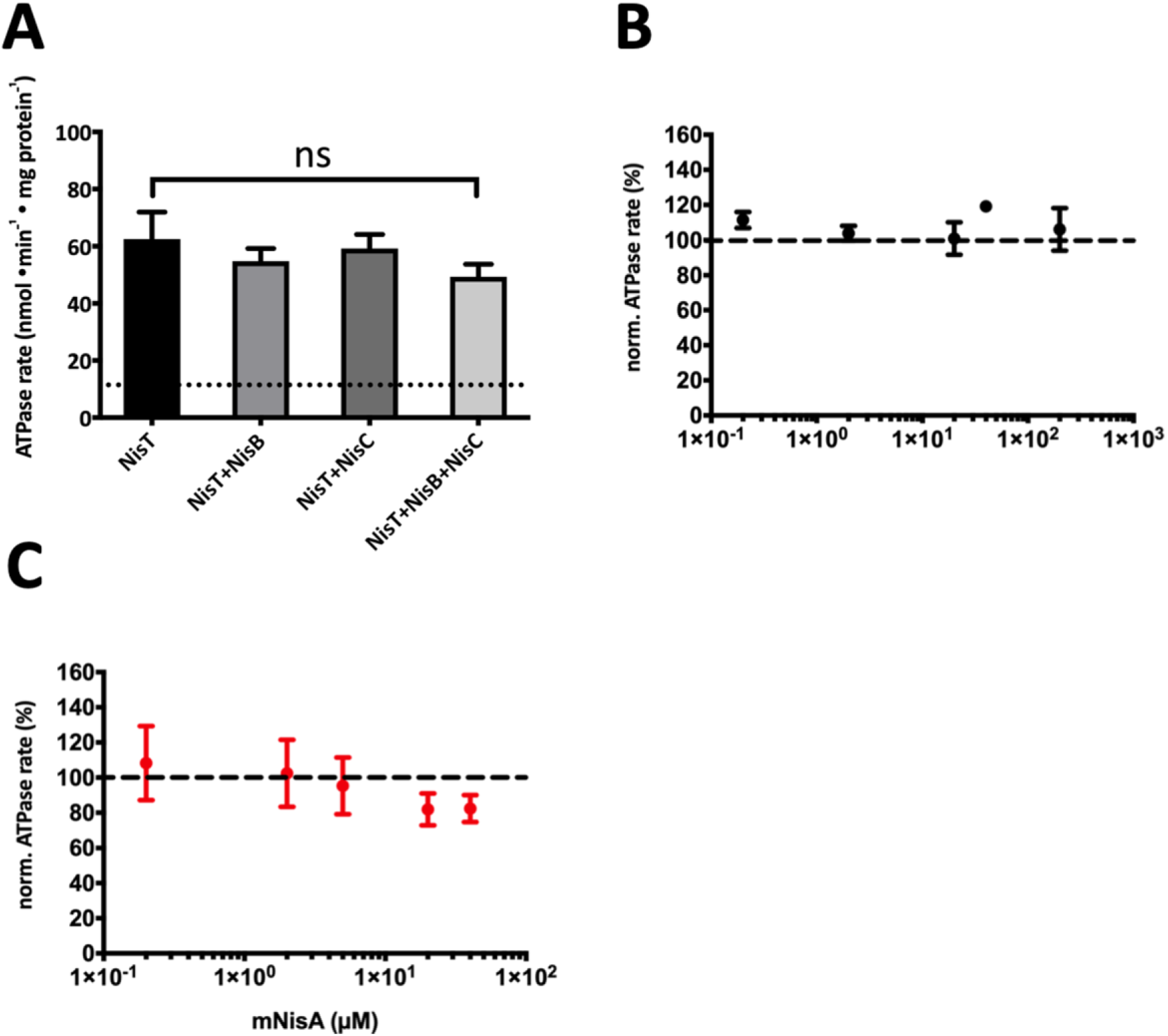
Influence of NisBC on the ATPase rate of NisT. The ATPase rate of purified 10HNisT was analyzed in the presence of the modification enzymes NisB and NisC, respectively. (**A**) ATPase rate of 10HNisT was plotted against variation of 10HNisT with the modification enzymes NisB, NisC and NisBC. It showed the normalized ATPase rate, in which the ATPase rate of 10HNisT was set to 100% (dashed line). (**B**) The substrate NisA_LP_ (black dots) and (**B**) mNisA (red dots) were used in the assay with 10HNisT in presence of NisBC. The normalized ATPase rate was plotted against various concentrations (µM), where the ATPase rate of 10HNisT and NisBC was set to 100% (dashed line). All assay assays were performed in at least four independent experiments and are represented as means ± SD (n=4). The means were analyzed by a one-way ANOVA. ns: not significant (p-value: ≥ 0.05)

Next, the ATPase rate of NisT with NisBC was investigated in presence of NisA_LP_ and mNisA, at concentrations ranging from 0 to 40 µM. The ATPase rate with the substrate NisA_LP_ was slightly increased but not in a concentration dependent manner (Figure 5B), while a decreasing effect on the ATPase rate was observed in presence of mNisA. Here, a concentration dependent manner was observed with a lowest value of 82% at 40 µM (Figure 5C).

### Interaction of NisT with NisBC

In 2017 the assembly of the nisin modification complex consisting of NisB_2_C and NisA was published and shed light on the stoichiometry and structure of the complex in solution (Reiners, Abts et al. 2017). Additionally, an influence of the last ring on complex disassembly was determined. The next step would be the interaction with NisT prior to secretion, as a proposed transient multimeric nisin modification/secretion complex (Siegers, Heinzmann et al. 1996), but detailed information about the interaction with the ABC transporter NisT are still missing.

Therefore, the interaction of NisT with NisB and NisC was investigated by a pull-down assay, in which 10HNisT was immobilized on NTA-magnetic beads (Qiagen). Initially, the interaction of NisT was tested with 10 µM NisB, NisC or NisBC. Interestingly, in all cases the interaction partner of NisT were observed in the elution fractions. This clearly shows a specific interaction of NisB and NisC with NisT independent of pre-NisA (Figure 6A). The controls, in which the NTA-magnetic beads were incubated with NisB and NisC without immobilized 10HNisT, revealed no protein in the elution fractions (Supplementary Figure 5).

**Figure 6:**
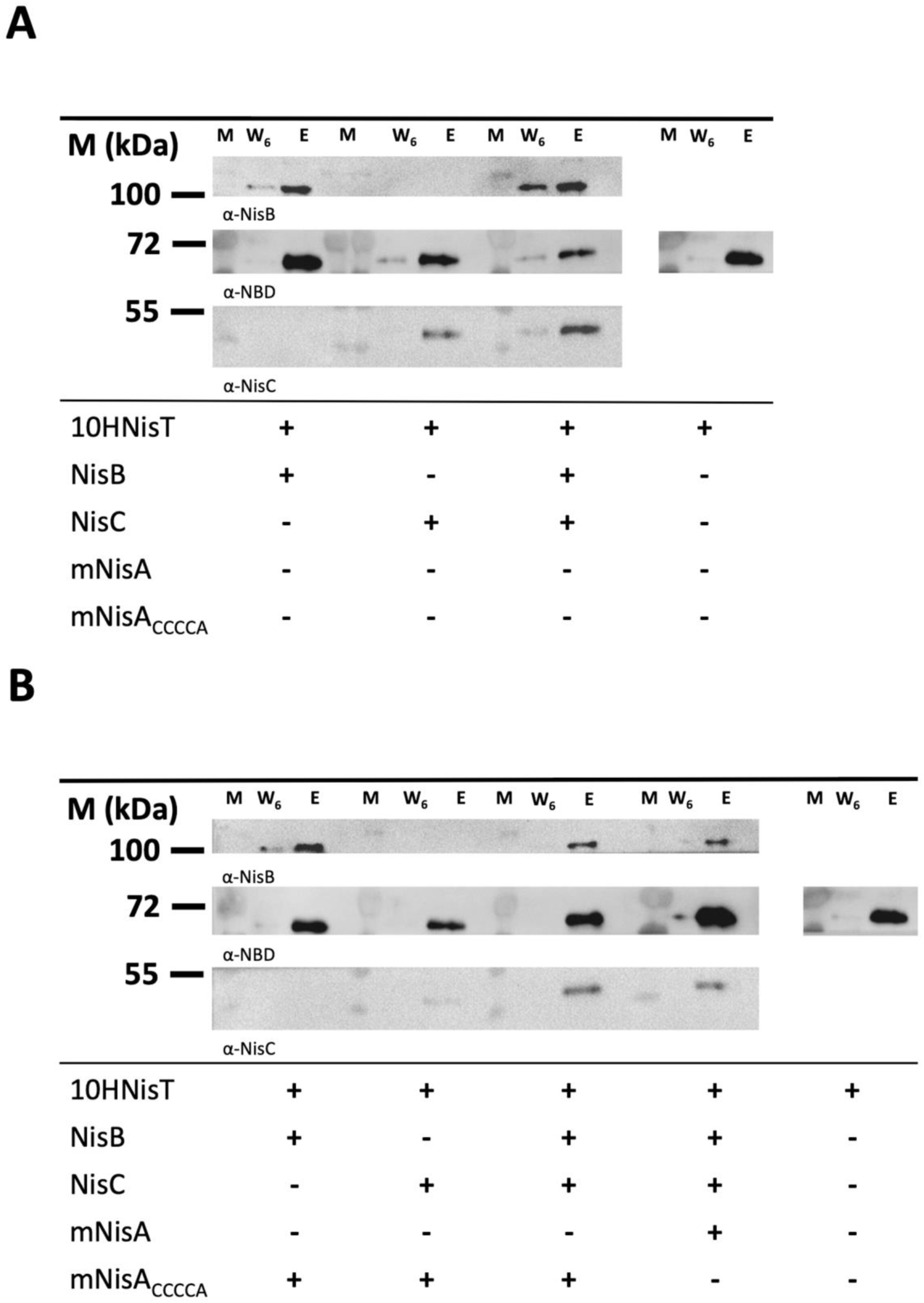
Pull-down assay of NisT with NisB and NisC. The interaction of 10HNisT with NisB and NisC was studied by a pull-down assay. The ABC transporter was immobilized on NTA-magnetic beads, the specific interaction partner were added and incubated. After six washing steps, the last washing step (W_6_) and the EDTA elution fraction (E) were analyzed by WB with the specific antibodies (α-NDB, α-NisB or α-NisC). Western blots displayed the eluted bands for NisB, NisT and NisC without substrate (**A**) and with substrates mNisA or mNisA_CCCCA_ (**B**). The pull-down assay was repeated three times and showed similar results. M: marker protein bands; +: protein was used in assay; −: protein was not used in assay

The same set up was used in the presence of the substrate mNisA_CCCCA_, which lacks the last lanthionine-ring (ring E) and stabilizes the maturation complex of NisBC (Reiners, Abts et al. 2017). This substrate showed no additional effect on the interaction of NisB with NisT. However, the interaction of NisT and NisC was affected and the amount of co-eluted NisC was strongly reduced (Figure 6B). After addition of NisB, the interaction of NisB and NisC with NisT was restored (Figure 6B). Noteworthy, the addition of mNisA instead of the ring-mutant shows an identical result on complex formation. Unfortunately, the analysis of all elution fractions with an antibody against the LP gave no signals for the substrates mNisA/mNisA_CCCCA_. This might be due to low concentrations of the peptides in the elution fractions.

In summary, this is the first time that beside the interaction of NisT and NisC, an interaction of NisT with NisB was shown. Even the co-elution of NisB and NisC with NisT as a complex in the presence of the substrates mNisA and the ring-mutant mNisA_CCCCA_ was observed. Although no enhancement of the interaction between NisT and NisBC in presence of substrates mNisA/ mNisA_CCCCA_ was detected, NisB plays a pivotal for complex stability.

## Discussion

The mechanism of lanthipeptide modifications was subject of many studies, but still only little is known about the secretion process of class I lanthipeptide ABC transporters (LanT). Only a few *in vivo* studies investigated the translocation of lanthipeptides nisin, subtilin, Pep5 or epidermin (Schnell, Engelke et al. 1992, Meyer, Bierbaum et al. 1995, Ra, Qiao et al. 1996, Izaguirre and Hansen 1997, Kuipers, de Boef et al. 2004). Amongst these systems, the nisin modification and secretion system (NisBTC) is the best studied one and is commonly employed to secrete nisin variants, lanthipeptides or non-lanthionine containing peptides (Kuipers, de Boef et al. 2004, Rink, Kuipers et al. 2005, van Heel, Kloosterman et al. 2016). It is commonly established, that nisin is ribosomally synthesized as a precursor peptide NisA and undergoes special post-translational modifications (e.g. Dha, Dhb, Lan and MeLan) (Newton, Abraham et al. 1953, Gross and Morell 1967, Ingram 1970). The PTM of the CP are installed in a coordinated manner by the nisin modification complex NisB_2_C (Khusainov, Heils et al. 2011, Reiners, Abts et al. 2017).

Not only the modification is a tightly coupled process, but also the secretion of mNisA by NisT. A proposed channeling mechanism through interaction with NisB/NisC might explain mNisA translocation. Here, mNisA can be detected in the medium within the first minute after induction (van den Berg van Saparoea, Bakkes et al. 2008). We characterized the LanT-type transporter NisT with respect to the secretion process and its specific activity. To study the mechanism of nisin secretion, we focused on two main topics: (I) *In vivo* secretion rate of NisA by NisT and (II) *in vitro* activity of NisT with and without substrate.

In 2008 van den Berg van Saparoea *et al*. conducted a kinetic analysis of nisin production with the strains NZ9700 and NZ9000 transformed with a two plasmid system (van den Berg van Saparoea, Bakkes et al. 2008). They demonstrated distinct contributions of the modification enzymes NisB and NisC with respect to lanthipeptide secretion and proposed that the secretion process of NisT occurred via a channeling mechanism. This hypothesis was supported by further *in vivo* studies, in which some mechanistic aspects of NisB and NisC modification were investigated (Lubelski, Khusainov et al. 2009, Khusainov, Heils et al. 2011, Khusainov and Kuipers 2013).

Although, these studies clearly demonstrated the dependence of pre-NisA secretion on the modification enzymes, it did not include a determination of the underlying kinetic parameters. To determine these kinetic parameters, we quantified the amount of secreted peptide via HPLC from different time points of various NZ9000 strains.

The first two kinetic parameters were V_max_ and K_0.5_, which were obtained by an allosteric sigmoidal analysis (Figure 2). Generally, our results are consistent with the aforementioned studies, in which strain NZ9000BTC had the highest V_max_ and the lowest K_0.5_ value reflecting a high secretion efficiency. The strains, which secreted uNisA (NZ9000T and NZ9000TC), show lower V_max_ and higher K_0.5_ values. Interestingly, we observed in our secretion assay some aberrations with respect to dNisA secretion. The expression of a catalytic-inactive NisC (H331A mutant) in the NisBTC system (strain NZ9000BTC_H331A_) did not restore the secretion level of dNisA to the WT level. This is in contrast to Lubelski *et al*., where a recovery of the pre-NisA secretion to WT level was observed (Lubelski, Khusainov et al. 2009). The secretion of dNisA by NZ9000BT has a higher V_max_ level and a lower K_0.5_ value. However, one has to consider the time scale of the secretion assays, which might explain this difference. In our assay, the early kinetics of secretion (also see (van den Berg van Saparoea, Bakkes et al. 2008)) might pronounce the differences between the strains more clearly as an end-point determination after an overnight secretion. The precise determination of secretion efficiency revealed a descending order of the pre-NisA secreting strains (NZ9000BTC > NZ9000BT > NZ9000BTC_H331A_ > NZ9000T > NZ9000TC). The secretion efficiency clearly shows, that mNisA is secreted at high rates by strain NZ9000BTC and every aberration of the secretion system reduced the rate at least by a factor of 2.2 (see strain NZ9000BT).

The third kinetic parameter was the apparent secretion rate V_S app._ (NisA•NisT^−1^•min^−1^), which we determined for the strains NZ9000BTC and NZ9000NisT, in which the secretion efficiency was the highest for NisBTC system (Figure 2). These values are in general an approximation to obtain a kinetic parameter, which can be related to other secretion system, e.g. the SecA translocon. In comparison to the nisin modification and secretion system (V_S app_: 191 ± 67 aa•s^−1^) the SecA translocon processes a secretion rate of 152-228 aa•s^−1^ per translocon (Robson, Gold et al. 2009). A lower secretion rate was determined for the hemolysin A type 1 secretion system (HlyA T1SS), which was approximately 16 aa•T1SS^−1^•s^−1^ (Lenders, Beer et al. 2016). Similar to the latter, we observed that ATP hydrolysis is essential for the secretion process as the H551A mutant (strains NZ9000BT_H551A_C) did not secrete mNisA. This mutant enables ATP binding and NBD dimerization but not ATP hydrolysis (Zaitseva, Jenewein et al. 2005).

As ATP hydrolysis is clearly important for ABC transporter mediated substrate translocation, we determined the *in vitro* activity of NisT in terms of ATPase rate without and with substrate. Here, the basal ATPase activity had a value of 79.9 ± 2.9 nmol•min^−1^•mg^−1^ with a K_m_ value of 0.37 ± 0.04 mM, which is in the range of other ABC transporter (Choudhury, Tong et al. 2014, Lin, Huang et al. 2015, Reimann, Poschmann et al. 2016, Bock, Zollmann et al. 2019) (Figure 3). In comparison to NisT, the Lan_C39P_T transporter NukT has a low V_max_ value of 12.6 nmol•min^−1^•mg^−1^, but it is stimulated by mNukA up to max. 500% (at 50 µM substrate). The cleaved substrate stimulates up to 200 % (at 25 µM substrate) and the unmodified substrate does not stimulate at all (Zheng, Nagao et al. 2017). In the case of NisT, we did not observed a substrate concentration dependent stimulation in the presence of NisA_LP_, uNisA, dNisA and mNisA (Figure 4). NisT is maximal stimulated to ∼ 140%. We extended the ATPase activity assay by addition of NisB and NisC to stimulate the WT system (NisBTC), in which the secretion of pre-NisA is the most efficient (Figure 5). The addition of Nis_LP_ had no effect on the ATPase rate. However, the addition of mNisA revealed an inhibiting effect on the ATPase rate with increasing substrate concentration (approximately 80% at 40 µM). Interestingly, a similar behavior was observed for PCAT1 (Lin, Huang et al. 2015). Another study on an ABC transporter homologue to PrtD from *Aquifex aeolicus* also noticed an inhibitory effect on ATPase activity after substrate addition (Morgan, Acheson et al. 2017). The open question is now, if NisT uses equivalent mechanism, in which the interaction of the modification/ secretion complex inhibits the ATPase rate prior to translocation.

In 1996 a multimeric enzyme complex of NisBTC was proposed, but the isolation of such a complex was not successful (Siegers, Heinzmann et al. 1996). Therefore, we choose to study the specific interaction of NisT with NisB and NisC via a pull-down assay. Such a pull-down assay was performed with His-tagged pre-NisA, where NisB and NisC were co-eluted from cytoplasmic fraction (Khusainov, Heils et al. 2011, Khusainov, Moll et al. 2013). In our study, we expanded this set up and used purified NisT, NisB, NisC and pre-NisA (Figure 6). We observed a specific interaction of NisT with NisC, which is in line with previously observed interaction of NisT with NisC via co-immunoprecipitation and yeast two-hybrid assay (Siegers, Heinzmann et al. 1996). Another specific interaction of the modification enzyme with the ABC transporter was shown for NukM and NukT. Here, the C-terminal domain of LanM (LanC-like domain; aa 480-917) interacts with TMD and NBD of NukT, but not with the C39 peptidase domain (Nagao, Aso et al. 2005). Besides the interaction of NisT with NisC, we also noticed an interaction of NisT with NisB, which was not observed in the above-mentioned study, but for SpaT and SpaB in a similar experiment (Kiesau, Eikmanns et al. 1997). Since secretion of dNisA by NZ9000BT was observed (Kuipers, de Boef et al. 2004), an interaction of NisT and NisB was also proposed. Furthermore, we observed for the first time the co-elution of NisBC with NisT in the proposed lanthionine synthetase complex. Remarkably, the co-elution of transporter with the modification enzymes is not increased by addition of the substrates mNisA or mNisA_CCCCA_. Similar amounts of the enzymes were co-eluted and we conclude that the interaction of NisT with NisB and NisC is independent of substrate. One exception was the addition of mNisA_CCCCA_ to NisT/NisC, in which the amount of co-elute NisC was reduced. Only the addition of NisB to the sample increased the co-elution of NisC. It is now commonly accepted that NisB represents the main component of the NisBTC modification/secretion complex (van den Berg van Saparoea, Bakkes et al. 2008, Lubelski, Khusainov et al. 2009) and based on our data, NisB stabilizes the NisBTC complex.

In summary, we have determined the kinetic parameter for the *in vivo* secretion of the nisin modification and secretion complex NisBTC. We demonstrated that alternations in the NisBTC system lead to impaired secretion of pre-NisA or even to no secretion, when an ATP hydrolysis deficient mutant of NisT was used. For an efficient secretion by NisT the modification enzymes NisB and NisC are prerequisite and their interaction with NisT enhances the secretion process by the proposed channeling mechanism (van den Berg van Saparoea, Bakkes et al. 2008). Interestingly, it is the function of NisC that is required, but the presence of NisB. Additionally, the *in vitro* activity data of NisT demonstrate that the lanthipeptide exporter is not stimulated by the interaction partner and substrate, respectively. The observed complex formation of NisB and NisC with NisT hint to the proposed lanthionine synthetase complex, in which NisT under goes a transition to a transport competent exporter in presence of only NisB, NisC and the substrate mNisA.

## Material and Methods

### Chemicals and Antibodies

Fos-choline 16 (FC-16) and CYMAL5 (C5) were obtained from Anatrace. Lyophilized nisin powder (2.5% nisin content) and insulin chain B were obtained from Sigma-Aldrich. The leader peptide of NisA was synthesized and obtained from JPT peptide technologies. Antibodies for NisT_NBD_, NisC and NisA_LP_ peptide were purchased from Davids Biotechnology (Germany) as polyclonal antibodies. NisB antibody was kindly provided by Dr. G. Moll (Lanthio Pharma; Groningen, Netherlands). All standard chemicals were purchased from Sigma-Aldrich or VWR.

### Bacterial strains and growth conditions

Strains of *Escherichia coli* and *Lactococcus lactis* and plasmids used in this study are listed in Supplementary File 1. The strains *E. coli* DH5α or BL21 were grown in LB medium at 37 °C under aerobic conditions with appropriate antibiotics (30 μg/ml kanamycin or 100 μg/ml ampicillin). The transformation of *E. coli* strains was performed following standard procedures.

The strain *L. lactis* NZ9000 (and its variants) was grown in M17 (Terzaghi and Sandine 1975) or minimal medium (MM) (Jensen and Hammer 1993, Rink, Kuipers et al. 2005) at 30 °C under semi aerobic conditions supplemented with 0.5% glucose (GM17/ GMM) and appropriate antibiotics (erythromycin or/and chloramphenicol at a final concentration of 5 μg/ml). To MM a vitamin mix (100x stock solution, 1x final) was added.

For transformation of *L. lactis* NZ9000 with the expression plasmids a standard procedure for preparation of competent cells and electroporation was used (Holo and Nes 1989).

### *In vivo* secretion assay

The supernatants of the *in vivo* secretion assay were analyzed by RP-HPLC. The secreted peptides were separated by an acetonitrile/water gradient after a 20% washing step on a C-18 RP-HPLC column. The eluted peptides were monitored via UV signal at 205 nm and collected fractions of 1 ml were collected and used for MALDI-TOF-MS analysis. The MS analysis confirmed the correct masses for pre-NisA peptides (uNisA, dNisA or mNisA) in fractions 30-36 min of the chromatogram (Supplementary Figure 1). The modified peptide mNisA with eight (−Met: 5689 Da) or seven (−Met: 5707 Da) dehydrations from strain NZ9000BTC was detected in fractions 34-36 min (Supplementary Figure 1A, Supplementary File 2). Furthermore, the modified peptides (dNisA) from strains NZ9000BTC_H331A_ and NZ9000BT with eight dehydrations (−Met: 5689 Da) were found in fractions 33-36 min (Supplementary Figure 1B/D, Supplementary File 2). In the case of the peptide from strains NZ9000T and NZ9000TC the retention time was shifted and the unmodified peptide was eluted in fractions 30-32 min (Supplementary Figure 1E/F). The corresponding molecular mass for unmodified pre-NisA (+Met: 5951 Da; −Met: 5834 Da) was verified (Supplementary File 2).

### Cloning of *nisT* and *nisT* variants

A nucleotide sequence for a MCS with 10H nucleotide sequence was ordered as a codon-optimized, synthetic gene fragment from Life Technologies to insert it into the pNZ-SV plasmid (AlKhatib, Lagedroste et al. 2014). The synthetic gene fragments were amplified by Phusion DNA polymerase (NEB) with the primer pair 10Hfor and 10Hrev (Supplementary File 4) for Gibson assembly. The plasmid pNZ-SV was amplified by Phusion DNA polymerase (NEB) with the primer pair infupNZ-SVfor and infupNZ-SVrev (Supplementary File 4) to linearize the vector. The gene fragment and the vector pNZ-SV were employed in the Gibson assembly by following the manufactures instructions (NEB). The Gibson assembly reactions were transformed into *E. coli* DH5α. The sequence of the construct pNZ-SV10H (Supplementary File 5) was verified by DNA sequencing (Microsynth Seqlab).

The *nisT* gene (accession number: Q03203) was amplified using genomic DNA from *L. lactis* NZ97000 (Kuipers, Beerthuyzen et al. 1993) as a template. Phusion DNA polymerase (NEB) with the primer pair infunisTor and infunisTrev (Supplementary File 4) was used to create overhang sequences for Gibson assembly. The plasmid pNZ-SV10H was amplified by Phusion DNA polymerase (NEB) with the primer pair linpNZ-SVfor and linpNZ-SVrev (Supplementary File 4) to linearize the vector. Subsequently, the gene and the linearized vector pNZ-SV10H were employed in the Gibson assembly and the reactions were transformed into *E. coli* DH5α. The sequence of the construct pNZ-SV10HnisT (Supplementary File 5) was verified by DNA sequencing (Microsynth Seqlab).

To generate the plasmid pIL-SVnisT the *nisT* gene from pNZ-SV10HnisT was amplified by Phusion DNA polymerase (NEB) with the primer pair infupIL-SVfor and infupIL-SVrev (Supplementary File 4) to create overhanging sequences. The plasmid pIL-SV (AlKhatib, Lagedroste et al. 2014) was linearized by Phusion DNA polymerase (NEB) with the primer pair termpNZfor and pnisArev (Supplementary File 4). The gene with overhang sequences and the vector was employed in Gibson assembly. The Gibson assembly reactions were transformed into *E. coli* DH5α. Additionally, the 10H nucleotide sequence was deleted by Phusion DNA polymerase (NEB) with the primer pair 10Hfor and infupNZ-SVrev (Supplementary File 4). The sequence of the construct pIL-SVnisT (Supplementary File 5) was verified by DNA sequencing (Microsynth Seqlab).

To generate the *nisT*_H551A_ mutant, a polymerase chain reaction using Pfu DNA polymerase (Thermo Fischer Scientific) or Pfu DNA polymerase (Promega), the template pNZ-SV10HnisT or pIL-SVnisT and the primer pair nisT_H551A_for and nisT_H551A_rev (Supplementary File 4) was performed according to standard procedures. The sequence of the constructs (Supplementary File 5) was verified by DNA sequencing (Microsynth Seqlab).

The plasmid pNZ-SVnisTNBD_H348_ was obtained by the deletion of the TMD sequence (1-347) from the plasmid pNV-SV10HnisT. The plasmid was amplified with Phusion DNA polymerase (NEB) with the primer pair ΔnisT_TMD_for and ΔnisT_TMD_rev (Supplementary File 4). The linear vector was ligated with T4-ligase (NEB) and transformed into *E. coli* DH5α. The sequence of the construct pNZ-SVnisTNBD_H348_ (Supplementary File 5) was verified by DNA sequencing (Microsynth Seqlab).

### Cloning of nisBTC and *nisBTC* variants

The plasmid pIL-SVnisBTC was generated from pIL-SV and pIL3BTC (Rink, Kuipers et al. 2005). The plasmid pIL3BTC was digested with the restriction enzymes NotI (NEB) and BstXI (NEB) to receive a fragment BTC containing the genes *nisB*, *nisT* and *nisC*. Next, pIL-SV (AlKhatib, Lagedroste et al. 2014) was also digested with NotI and BstXI (pIL-SV**). The fragment BTC and pIL-SV** were ligated with T4-ligase (NEB) and transformed into *E. coli* DH5α. The sequence of the construct pIL-SVnisBTC (Supplementary File 5) was verified by DNA sequencing (Microsynth Seqlab).

By using Phusion DNA polymerase (NEB) with the appropriate primer pairs (Supplementary File 4) the gene deletions of *nisB*, *nisC* or *nisT* were performed to generate pIL-SVnisBTC derivatives. Subsequently, the linearized vectors were ligated with T4-ligase (NEB) and transformed into *E. coli* DH5α. The sequence of the constructs (Supplementary File 5) were verified by DNA sequencing (Microsynth Seqlab).

To generate *nisT*_H551A_ and nisC_H331A_ mutants, a polymerase chain reaction using PfuUltra II Fusion DNA polymerase (Agilent Technologies), the template pIL-SVnisBTC and the appropriate pair of oligonucleotides (Supplementary File 4) was performed according to standard procedures. The sequence of the new constructs (Supplementary File 5) were verified by DNA sequencing (Microsynth Seqlab).

### *In vivo* secretion assay: Expression and secretion of pre-NisA

Strain *L. lactis* NZ9000 harboring the plasmids pIL-SVnisBTC and pNZ-SVnisA hereupon termed NZ9000BTC (Supplementary File 1) was used to investigate the *in vivo* secretion activity of the nisin modification and secretion system (NisBTC). The use of pIL-SVnisBTC derivatives and pNZ-SVnisA for transformation into *L. lactis* NZ9000 led to strains described in detail in Supplementary File 1. For each secretion experiment new transformants were prepared and used to inoculate GM17 (Erm+Cm) with one colony. The overnight culture was centrifuged at 4000xg for 20 min and cells were resuspended in GMM. Subsequently, 0.5 l GMM (Erm+Cm) were inoculated to OD_600_ of 0.3 and incubated at 30°C. After 60-90 min the culture (OD_600_ of 0.4-0.5) was induced with 10 ng/ml nisin (powder from Sigma-Aldrich dissolved in 50 mM lactic acid). A 50 ml sample before induction (0 hour) and every other hour (1-6h) was taken. For each sample the cell were harvested by centrifugation at 4000xg for 20 min. Subsequently, the cells were resuspended in R-buffer (50 mM Na-Phosphate buffer, pH 8, 100 mM KCl, 20% glycerol) to an OD_600_ of 200, flash frozen in liquid nitrogen (N_2_) and stored at −80 °C until further use. The supernatant was additionally centrifuged at 17,000xg for 20 min at 8°C. Supernatants were kept on ice before the RP-HPLC analysis. Furthermore, 2 ml or 10 ml of the supernatant were precipitated by 1/10 volume (10%) TCA. TCA samples were incubated at 8°C overnight. The TCA-precipitated peptide was centrifuged at 17,000xg for 20 min at 8°C and consecutively washed three-times with ice-cold acetone. The pellets were vacuum-dried and resuspended in 60 µl per OD_600_ 1 of 1x SDS-PAGE loading dye containing 5 mM β-mercaptoethanol (β-ME). These resuspended TCA-pellets were analyzed by Tricine-SDS-PAGE and Western blot.

### *In vivo* secretion assay: Analysis of cell pellets

The resuspened cell pellets were thawed on ice and 1/3 (w/v) glass beads (0.3 mm diameter) were added. Cells were disrupted on a vortex-shaker (Disrutor Genie, Scientific Industries). A cycle of 2 min disruption and 1 min incubation on ice was repeated five times. A low spin step at 17,000xg for 30 min at 8°C and subsequently a high spin step at 100,000xg for 120 min at 8°C was performed. The supernatant of the latter centrifugation step represents the cytoplasmic fraction and the pellet corresponds to the membrane fraction. The SDS-PAGE samples of cytoplasmic and membrane fractions were prepared by adding 4x SDS-PAGE loading dye containing 5 mM β-ME and used for SDS-PAGE as well as Western blot analysis.

### *In vivo* secretion assay: Analysis of culture supernatant

The culture supernatants containing pre-NisA variants were analyzed by RP-HPLC (Agilent Technologies 1260 Infinity II). A LiChrospher WP 300 RP-18 end-capped column and an acetonitrile/water solvent system were used as described previously (Abts, Montalban-Lopez et al. 2013). In the case of modified pre-NisA 300 µl and for all other variants (dehydrated/unmodified) 500 µl sample were injected. Prior to the gradient (20-50% acetonitrile) a washing step of 20% acetonitrile was used to remove most of the casein peptides. The peptide amount or pre-NisA in the supernatant was determined using the peak area integration analyzed with the Agilent Lab Advisor software. A calibration with known amounts of nisin or insulin chain B was used to obtain a linear regression line. Unknown amounts in the *in vivo* secretion assay samples were calculated as nmol or µM based on this linear regression line.

### *In vivo* secretion assay: Determination of kinetic parameter

The amount of secreted pre-NisA of the different *L. lactis* NZ9000 strains were plotted against time and fitted using an allosteric sigmoidal fit (1). Note that y is the amount secreted peptide (nmol), V_max_ the maximal secreted amount, x is time (min), K_0.5_ the time point at which 50% of V_max_ is present and h is the Hill slope indicating cooperatively. The analysis was performed using Prism 7.0c (GraphPad).

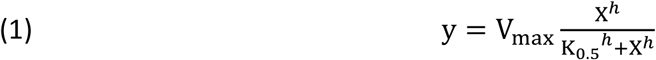

The apparent secretion rate (V_s app_) was determined by plotting the amount of NisA and NisT against time (min). The values were fitted using a linear regression (2). Note that y is the amount of NisA molecules per NisT molecules (NisA•NisT^−1^), m is the slope V_s app_ (NisA•NisT^−1^•min^−1^), x is the time (min), and b the y –axis interception. The analysis was performed using Prism 7.0c (GraphPad).

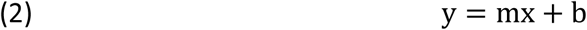

### Expression and purification of NisT

*L. lactis* NZ9000 strain was transformed with pNZ-SV10HnisT and placed on SMGG17 agar plates containing 5 µg/ml erythromycin. A GM17 (Erm) overnight culture was inoculated with one colony and incubated at 30°C. A GM17 (Erm) main culture was inoculated to an OD_600_ of 0.1 with the overnight culture. After 3 h incubation, expression was induced by adding 10 ng/ml nisin (powder from Sigma-Aldrich dissolved in 50 mM lactic acid) and further grown for additional 3 h. Cells were harvested by centrifugation at 4000xg for 20 min at 8°C and resuspened in R-buffer (50 mM Na-Phosphate buffer, pH 8, 100 mM KCl, 20% glycerol) to an OD_600_ of 200. To the resuspened cells 10 mg/ml lysozyme was added and incubated at 30°C for 30 min. Prior to cell disruption, cells were incubated on ice for 15min. The cell suspension was passed through a homogenizer (M-110P, Microfluidics System) at 1.5 kbar at least four times. The homogenized cell suspension was centrifuged at 12,000xg for 30 min at 8°C. Subsequently, the supernatant was centrifuged at 100,000xg for 120 min at 4°C to collect the membrane fraction. Membranes were resuspended with R-buffer containing 10 mM imidazole and 0.5 mM AEBSF. The total membrane protein concentration was measured by BCA assay (Thermo Fischer Scientific) and the concentration was adjusted to 5-7.5 mg/ml. Membranes were solubilized with 1% (w/v) of the lipid-like detergents FC-16 (Anatrace) for 1 h at 8°C. Insoluble material was removed by centrifugation at 100,000xg for 30 min at 4°C. The supernatant was applied to a 5ml IMAC (Immobilized Metal-Ion-Affinity Chromatography) HiTrap Chelating column (GE Healthcare) preloaded with 100 mM zinc sulphate and equilibrated with low IMAC1 buffer (50 mM Na-Phosphate buffer, pH 8, 100 mM KCl, 20% glycerol and 10 mM imidazole) containing 0.5 mM AEBSF and 0.005% FC-16. Consecutively, non-bound protein was washed and the buffer was exchanged to low IMAC2 buffer (50 mM Tris-HCl pH 8, 100 mM KCl, 10% glycerol, 10 mM imidazole, 0.5 mM AEBSF and 0.005% FC-16). After an additional washing step with 30% high IMAC buffer (50 mM Tris-HCl pH 8, 100 mM KCl, 10% glycerol, 150 mM histidine, 0.5 mM AEBSF and 0.005% FC-16), 10HNisT was eluted with 100% high IMAC buffer. The elution fractions containing NisT were pooled, 10 mM DTT was added and further concentrated with a Vivaspin20 100 kDa molecular weight cut off (MWCO) centrifugal concentrator (Sartorius AG). Next, a size exclusion chromatography (SEC) was performed, where the concentrated protein sample was applied onto a Superose 6 10/300 GL column (GE Healthcare) equilibrated with SEC buffer (25 mM Tris-HCl pH 8, 50 mM KCl, 10% glycerol, 0.5 mM AEBSF, 2 mM DTT and 0.0015% FC-16). The main peak fractions were analyzed via SDS-PAGE and further concentrated via a Vivaspin6 100 kDa MWCO centrifugal concentrator (Sartorius AG) until a concentration of 50 µM was reached. The protein concentration was determined by NanoDrop spectrophotometer (Thermo Fischer Scientific) using a molar extinction coefficient of 86,180 M^−1^•cm^−1^ and the molecular mass of 72.6 kDa. Aliquots of 50 µM 10HNisT were flash frozen in liquid N_2_ and stored at −80 °C until further use. The NisT variants 10HNisT_H551A_ was expressed following the same protocol.

### Expression and purification of NisT_NBD_

*L. lactis* NZ9000 was transformed with pNZ-SVnisTNBD_H348_ and placed on SMGG17 agar plates containing 5 µg/ml erythromycin. A GM17 (Erm) overnight culture was inoculated with one colony and incubated at 30°C. A GM17 (Erm) main culture was inoculated to an OD_600_ of 0.1 with the overnight culture. After 90 min incubation the culture was incubated on ice. Then expression was induced by adding 10 ng/ml nisin (powder from Sigma-Aldrich dissolved in 50 mM lactic acid) and further grown for additional 3 h at 20 °C. Cells were harvested by centrifugation at 4000xg for 20 min at 8°C and resuspened in R-buffer (50 mM Na-Phosphate buffer, pH 8, 100 mM KCl, 20% glycerol) to an OD_600_ of 200. Then 10 mg/ml lysozyme was added and incubated at 30°C for 30 min. Prior to cell disruption, cells were incubated on ice for 15min. Then the cell suspension was passed through a homogenizer (M-110P, Microfluidics System) at 1.5 kbar at least four times. The homogenized cell suspension was centrifuged at 12,000xg for 30 min at 8°C. Subsequently, the supernatant was centrifuged at 100,000xg for 120 min at 4°C. The supernatant was applied to a 5ml IMAC HiTrap Chelating column (GE Healthcare) preloaded with 100 mM zinc sulphate and equilibrated with low IMAC1 buffer (50 mM Na-Phosphate buffer, pH 8, 100 mM KCl, 20% glycerol and 10 mM imidazole). Consecutively, non-bound protein was washed by a 30% step with high IMAC buffer (50 mM Na-Phosphate buffer, pH 8, 100 mM KCl, 20% glycerol, 500 mM imidazole). The protein 10HNisT_NBDH348_ was eluted by a gradient of 30 to 100% high IMAC buffer. The elution fractions containing NisT_NBDH348_ (T_NBD_) were pooled, 10 mM DTT was added and further concentrated with an Amicon 10 kDa MWCO centrifugal concentrator (Millipore). Next, the concentrated protein sample was applied onto a HiLoad Superdex 200 16/60 prep grade column (GE Healthcare) equilibrated with SEC buffer (25 mM CAPS pH 10, 20% glycerol, 2 mM DTT). The main peak fractions are analyzed via SDS-PAGE and further concentrated with an Amicon 10 kDa MWCO centrifugal concentrator (Millipore). The protein concentration was determined by NanoDrop spectrophotometer (Thermo Fischer Scientific) using a molar extinction coefficient of 45,840 M^−1^•cm^−1^ and the molecular mass of 32.2 kDa. Aliquots of 30 µM T_NBD_ were flash frozen in liquid N_2_ and stored at −80 °C until further use.

### Expression and purification of nisin modification enzymes NisB and NisC

NisB and NisC were expressed and purified as described previously (Mavaro, Abts et al. 2011, Abts, Montalban-Lopez et al. 2013, Reiners, Abts et al. 2017). Aliquots of concentrated proteins (90 µM (NisB) or 110 µM (NisC)) were flash frozen in liquid nitrogen and stored at −80 °C until further use. For the *in vitro* ATPase activity assay and pull-down assay the buffer of the proteins was exchanged via a PD SpinTrap G-25 spin columns (GE Healthcare) to activity assay buffer (25mM Tris-HCl pH 7.5, 50 mM KCl) containing 400 mM glutamate and 0.4% CYMAL5 (3 x cmc). Proteins were stored on ice and directly used for the *in vitro* assays. The protein concentration was determined by NanoDrop spectrophotometer (Thermo Fischer Scientific) by using the theoretical molar extinction coefficient and the molecular mass of the proteins.

### Expression and purification of NisA variants

All pre-NisA variants used in this study were expressed and purified as described previously with modifications (Lubelski, Khusainov et al. 2009, Mavaro, Abts et al. 2011). Briefly, pre-NisA variants were purified after 5 h expression in MM from 2 l (0.5 l in the case of the secretion experiments) culture supernatant via cation-exchange chromatography (cIEX). The cell-free supernatant was diluted 1:1 with LA buffer (50 mM lactic acid, pH 3) and applied to a 5 ml HiTrap SP Sepharose column (GE Healthcare). The column was washed by applying a pH gradient from 100 % LA buffer to 100% H buffer (50 mM HEPES-NaOH, pH 7). Finally, the peptides were eluted with H buffer containing 1 M NaCl. The fractions containing the peptides were pooled and concentrated with an Amicon (Millipore) centrifugal concentrator. First, the flow through (FT) fraction of a 30 kDa molecular weight cut off (MWCO) concentrator was collect. Then a 3 kDa MWCO filter was used to concentrate the peptides of the FT fraction. Aliquots of concentrated peptides were flash frozen in liquid N_2_ and stored at −80 °C until further use. For concentration determination the pre-NisA variants were analyzed by RP-HPLC (Agilent Technologies 1260 Infinity II) with a LiChrospher WP 300 RP-18 end-capped column and an acetonitrile/water solvent system as described previously (Abts, Montalban-Lopez et al. 2013).

### *In vitro* ATPase activity assay

The ATPase activity of NisT was determined with the malachite green assay as described previously with experimental alterations (Infed, Hanekop et al. 2011). In this assay the release of inorganic orthophosphate after ATP hydrolysis was colorimetric quantified based on a Na_2_HPO_4_ standard curve.

All reactions were performed at 30°C in a total volume of 30 μl in activity assay buffer containing 0.4% CYMAL5 and 10 mM MgCl_2_.

In each reaction ∼2 μg of detergent-solubilized and purified NisT was used and the reaction was started by adding ATP (0-5 mM). The background of the reaction was a sample without MgCl_2_. After 30 min the reaction was stopped by transferring 25 μl of each reaction into a 96-well plate containing 175 μl stop-solution (20 mM sulphuric acid). Consecutively, 50 μL of a staining solution (0.096% (w/v) malachite green, 1.48% (w/v) ammonium heptamolybdate and 0.173% (w/v) Tween-20 in 2.36 M sulphuric acid) was added. After 10 min the amount of free inorganic orthophosphate was quantified by measuring the absorption at 595 nm using an iMark microplate reader (Bio-Rad).

The specific ATPase activity of NisT was plotted against ATP concentrations and fitted using the Michaelis−Menten equation (3). Note that y is the reaction velocity, V_max_ the maximal reaction velocity, x is the substrate concentration and K_m_ the Michalis-Menten constant. The analysis was performed using Prism 7.0c (GraphPad).

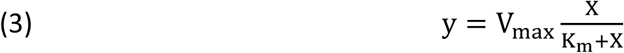

For the reactions with substrates (pre-NisA variants) or interaction partners (NisB and NisC) NisT was pre-incubated at 30°C for 10 min before ATP was added to start the reaction. All reactions were performed at 30°C in a total volume of 30 μl in activity assay buffer containing 0.4% CYMAL5, 400 mM glutamate and 10 mM MgCl_2_. In each reaction ∼2 μg of detergent-solubilized and purified NisT was used and the reaction was started by adding 5 mM ATP and stopped after 15 min following the procedure described above. In this reaction the concentration of the different substrates (0-40 µM) and /or interaction partner was varied and the ATPase activity was normalized to the specific ATPase activity of NisT without substrate/interaction partner. In these cases, the background was subtracted prior to normalization.

### *In vitro* pull-down assay

The immobilization of 10HNisT to Ni-NTA magnetic beads (Quiagen) was performed as described in the manufacture’s manual. In brief, ∼ 15 µg 10HNisT was incubated with Ni-NTA magnetic beads for 30 min at 30°C. Excess of protein was removed by three washing steps with activity assay buffer containing 0.4% CYMAL5 and 400 mM glutamate. Interaction partners NisC and NisB were incubated in 1:1 molar ratio (but > 10x molar excess to NisT) separately with or without mNisA/ mNisA_CCCCA_ (20x molar excess to NisT) in activity assay buffer containing 0.4% CYMAL5, 400 mM glutamate and 5mM MgATP for 15 min on ice. Next, interaction partner were added to 10HNisT immobilized to Ni-NTA magnetic beads and incubated for 1 h at 30°C. Positive control (only 10HNisT) and negative control (NisB, NisC) samples were prepared by incubating the proteins with Ni-NTA magnetic beads separately. After binding the Ni-NTA magnetic beads were washed six times with activity assay buffer. Finally, 10HNisT was eluted by adding activity assay buffer containing 50 mM EDTA. The SDS-PAGE samples of pull-down assay fractions were prepared by adding 4x SDS-PAGE loading dye containing 5 mM β-ME and used for Western blot analysis.

### SDS-PAGE and immunoblotting for protein/ peptide analysis

In general the sodium dodecyl sulfate−polyacrylamide gel electrophoresis (SDS-PAGE) experiments were performed using standard procedures (Laemmli 1970). In the SDS-PAGE gels the acrylamide portion was 10% to have a separation range from 30 to 120 kDa for the proteins NisC (∼48 kDa), NisT (∼69 kDa) and NisB (∼117 kDa).

All peptides (e.g. pre-NisA variants) from secretion experiments or from cIEX purification were analyzed by Tricine-SDS−PAGE (Schagger 2006). For Tricine-SDS-PAGE gels (12%) a Mini-Protean system (Bio-Rad) was used. Tricine-SDS−PAGE and SDS−PAGE gels were stained with colloidal coomassie (cc) (Dyballa and Metzger 2009).

All immunoblotting experiments were conducted following standard procedures. For the quantification of NisT in the membrane fractions (*in vivo* secretion assay) various amounts of a T_NBD_ standard (stock solution 12.5 µg/ml) was added to create a calibration curve. The band intensities on the Western blots were processed and determined by ImageJ (Schneider, Rasband et al. 2012). Then the amount of NisT was determined as pmol protein of the different membrane fractions of the time points 2-6 h.

### Mass spectrometry analysis

Pre-NisA variants were either desalted via ZipTip (C18 resin) purification accordingly to the manufacture manual (Merk-Millipore) or by RP-HPLC and vacuum dried.

For the MALDI-TOF-MS analysis the vacuum dried pellet was dissolved in 50% acetonitrile solution containing 0.1% TFA and analyzed as described elsewhere (Lagedroste, Reiners et al. 2019).

## Acknowledgements

We thank Peter Tommes for support and analysis of MALDI-TOF-MS experiments. We are greatly obliged to Diana Kleinschrodt and Iris Fey (former Protein Production Facility of HH) for their support of cloning the constructs. We also thank Olivia Spitz, Ioannis Panetas and Didem Kaya for their support during the early stages of the project. We are indebted to all members of the Institute of Biochemistry for stimulating discussions and support during the project.

## Additional information

### Notes

The Authors declare no competing financial interests.

## Author contributions

M.L., S.H.J.S. and L.S. designed the experiments, M.L. performed the experiments, J.R. purified NisB and NisC, M.L., S.H.J.S. and L.S. analyzed and interpreted the data, M.L., S.H.J.S. and L.S. wrote and revised the manuscript.

## Figures

**Supplementary Figure 1:**
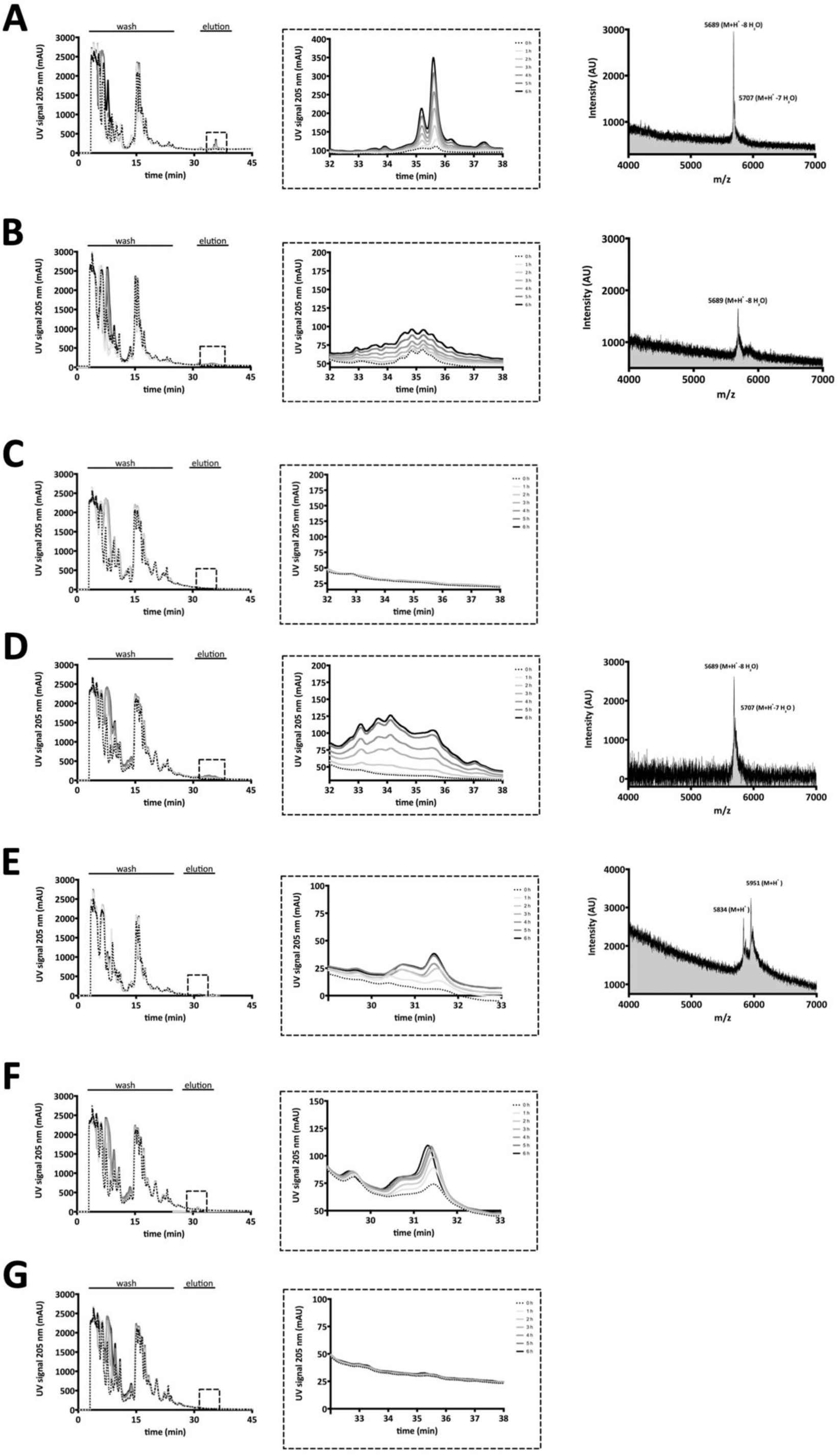
RP-HPLC chromatograms and MALDI-TOF-MS spectra of *in vivo* secretion assay. The supernatants of pre-NisA secreting *L. lactis* NZ9000 strains were employed for an RP-HPLC analysis. The pre-NisA variants (mNisA, dNisA and uNisA) were separated from other peptides in the supernatant by an acetonitrile/water gradient on a C-18 RP-HPLC column (left panel). The elution fractions (dashed square; middle panel) were further analyzed by MALDI-TOF-MS (right panel) to verify the correct mass. Integration of the corresponding peaks enables the determination of peptide amounts (nmol). Supernatant analysis of *L. lactis* strains (**A**) NZ9000BTC, (**B**) NZ9000BTC_H331A_, (**C**) NZ9000BT_H551A_C, (**D**) NZ9000BT, (**E**) NZ9000T, (**F**) NZ9000TC and (**G**) NZ9000BC.

**Supplementary Figure 2:**
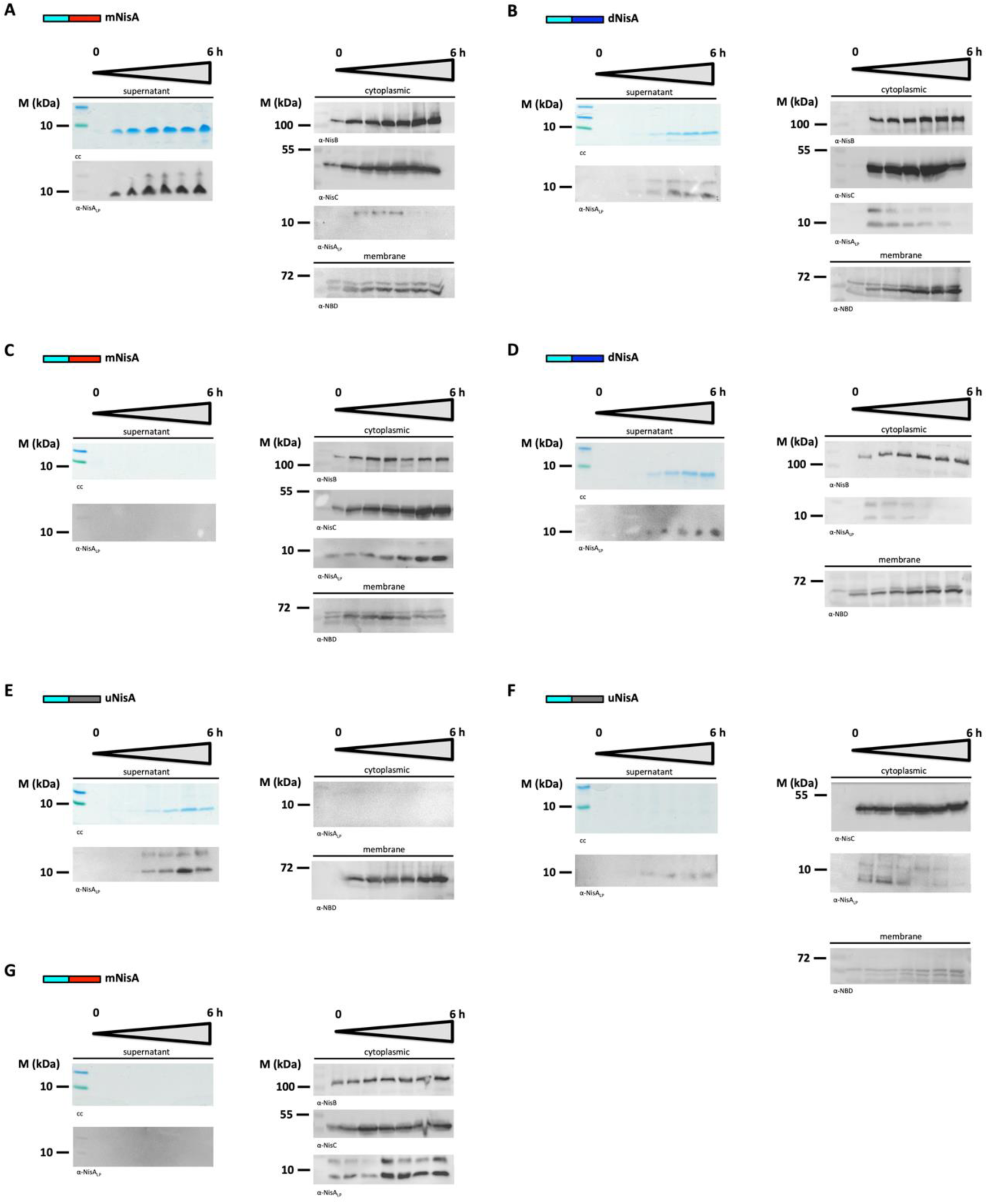
Tricine-SDS-PAGE and Western blot analysis of *in vivo* secretion assay. The supernatants of pre-NisA secreting *L. lactis* NZ9000 strains from different time points were TCA-precipitated and the pellets were analyzed by Tricine-SDS-PAGE and Western blot using an α-NisA_LP_ antibody (left panel). The Tricine-SDS-PAGE gels were stained by colloidal Coomassie (cc). The cytoplasmic and membrane fractions of the cell from different time points were analyzed by Western blot (right panel) with the specific antibodies (α-NisB, α-NisA_LP_, α-NisC and α-NBD). Sample analysis of *L. lactis* strains **A**) NZ9000BTC, (**B**) NZ9000BTC_H331A_, (**C**) NZ9000BT_H551A_C, (**D**) NZ9000BT, (**E**) NZ9000T, (**F**) NZ9000TC and (**G**) NZ9000BC. M: marker protein bands

**Supplementary Figure 3:**
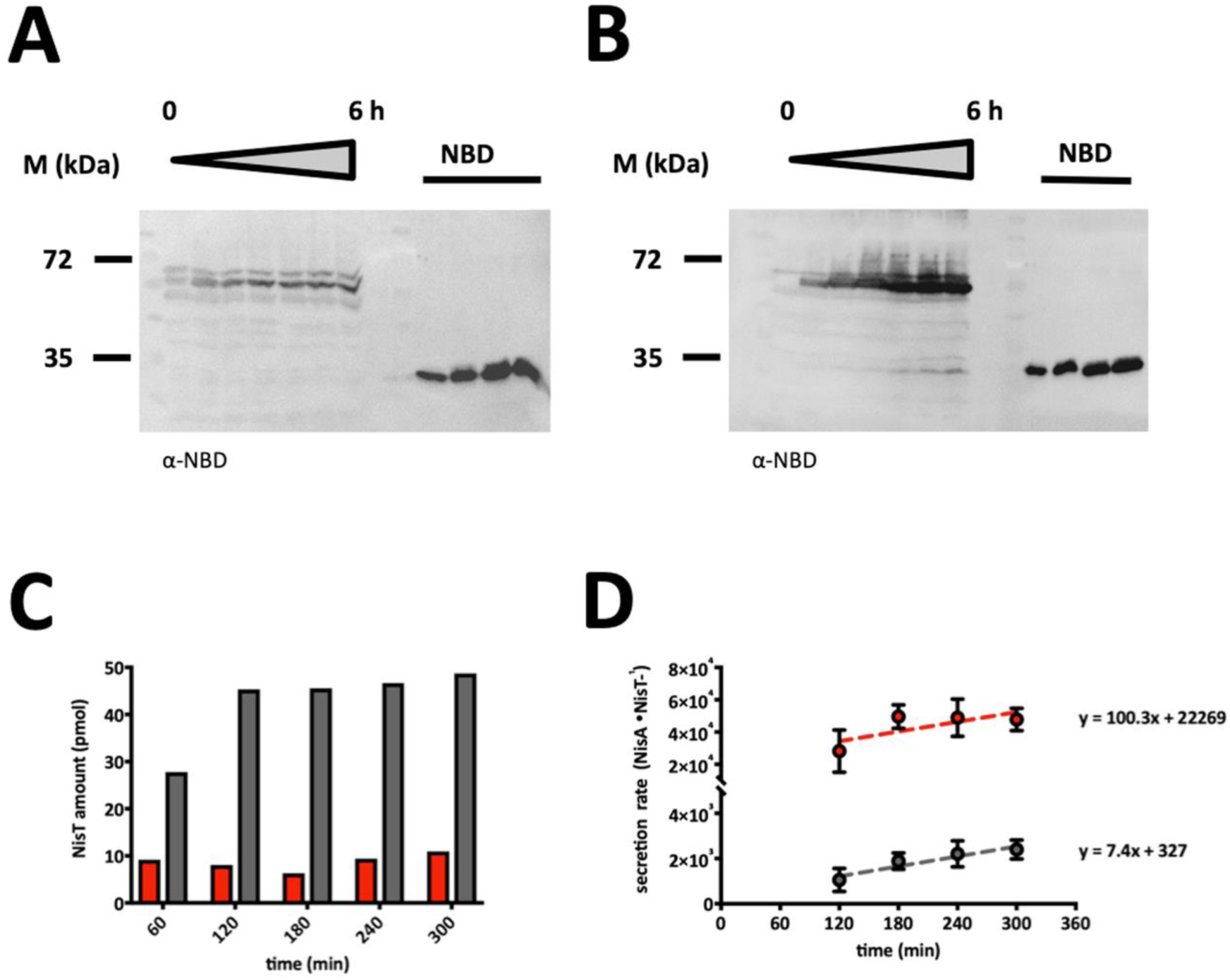
Determination of NisT amount of *in vivo* secretion assay samples. Western blot analysis of membrane fractions from strains (**A**) NZ9000BTC, (**B**) NZ9000BT and (**C**) NZ9000T with an antibody raised against the NBD (α-NBD). Standard of known amount of T_NBD_ was used to determine the (**D**) amount of NisT in the membrane (pmol) for time points 2-5 h. (**E**) A linear regression of plotted values of nmol NisA per nmol NisT against the time (min) results in a apparent secretion rate (V_s app_). M: marker protein bands

**Supplementary Figure 4:**
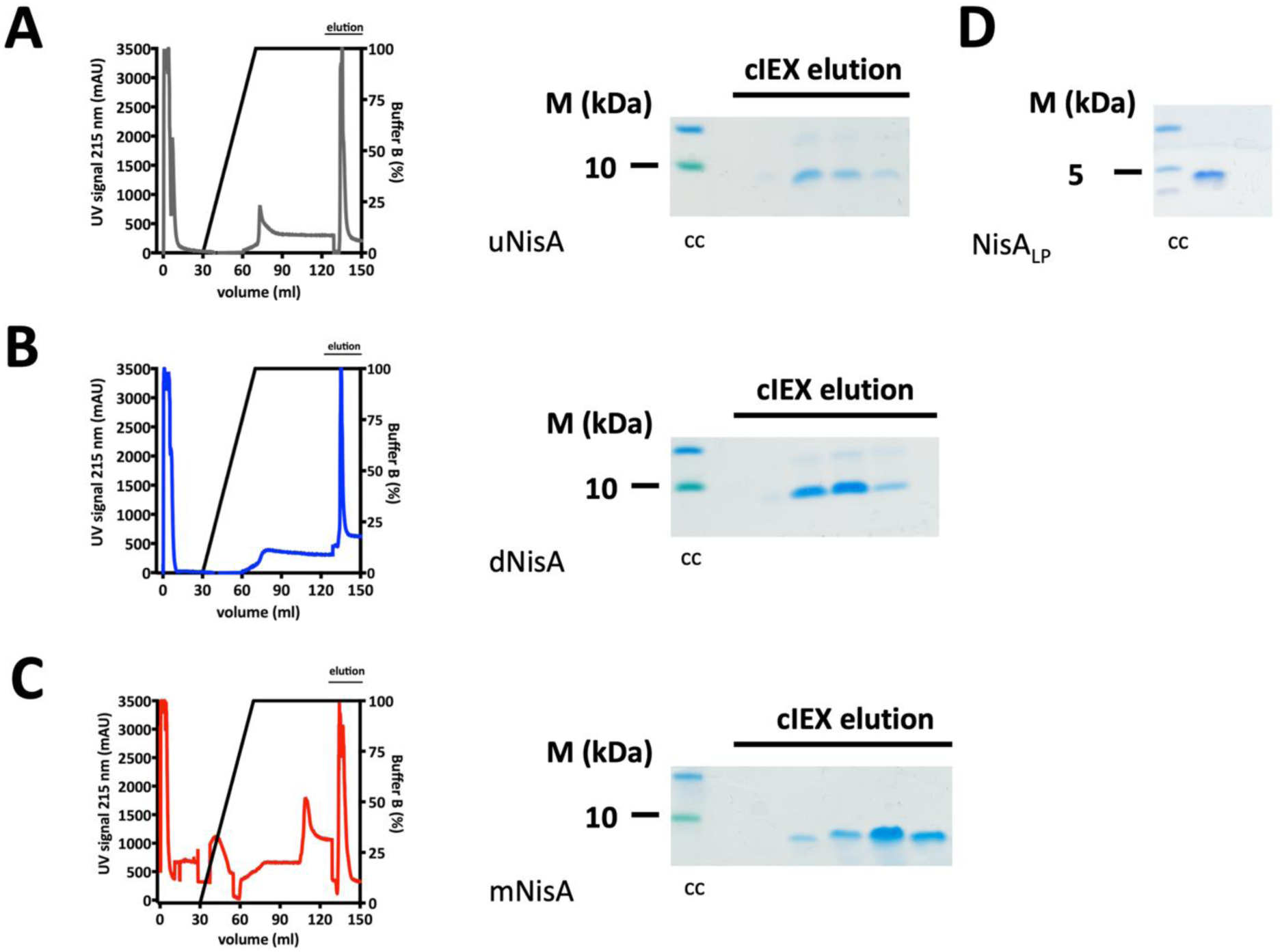
Chromatograms of cIEX from uNisA, dNIsA and mNisA. The cIEX chromatograms and Tricine-SDS-PAGE of pre-NisA variants (**A**) uNisA (grey), (B) dNisA (blue) and (C) mNisA (red). (**D**) The purity of the NisA leader peptide (NisA_LP_) was controlled by Tricine-SDS-PAGE. The Tricine-SDS-PAGE gels were stained by colloidal Coomassie (cc). M: marker protein bands

**Supplementary Figure 5:**
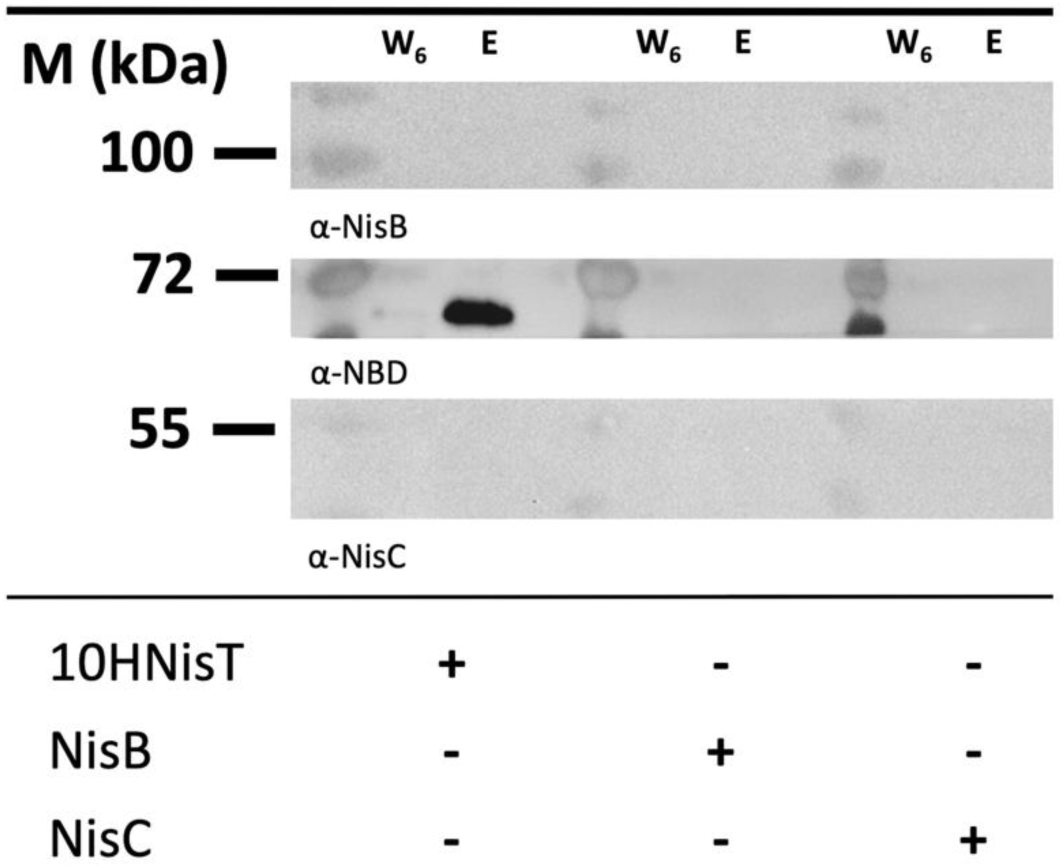
Western blot of pull-down assay with 10HNisT, NisB and NisC. Controls of the pull-down assay were blotted and analyzed with the specific antibodies of α-NisB, α-NBD and α-NisC. Samples of the sixth washing step (W_6_) and the elution fractions (E) were used to detect 10HNisT, NisB or NisC in the samples. M: marker protein bands; +: addition of protein; −: no protein added

## References

Abts, A., M. Montalban-Lopez, O. P. Kuipers, S. H. Smits and L. Schmitt (2013). “NisC binds the FxLx motif of the nisin leader peptide.” Biochemistry 52(32): 5387–5395.

AlKhatib, Z., M. Lagedroste, I. Fey, D. Kleinschrodt, A. Abts and S. H. Smits (2014). “Lantibiotic immunity: inhibition of nisin mediated pore formation by NisI.” PLoS One 9(7): e102246.

AlKhatib, Z., M. Lagedroste, J. Zaschke, M. Wagner, A. Abts, I. Fey, D. Kleinschrodt and S. H. Smits (2014). “The C-terminus of nisin is important for the ABC transporter NisFEG to confer immunity in Lactococcus lactis.” Microbiologyopen 3(5): 752–763.

Arnison, P. G., M. J. Bibb, G. Bierbaum, A. A. Bowers, T. S. Bugni, G. Bulaj, J. A. Camarero, D. J. Campopiano, G. L. Challis, J. Clardy, P. D. Cotter, D. J. Craik, M. Dawson, E. Dittmann, S. Donadio, P. C. Dorrestein, K. D. Entian, M. A. Fischbach, J. S. Garavelli, U. Goransson, C. W. Gruber, D. H. Haft, T. K. Hemscheidt, C. Hertweck, C. Hill, A. R. Horswill, M. Jaspars, W. L. Kelly, J. P. Klinman, O. P. Kuipers, A. J. Link, W. Liu, M. A. Marahiel, D. A. Mitchell, G. N. Moll, B. S. Moore, R. Muller, S. K. Nair, I. F. Nes, G. E. Norris, B. M. Olivera, H. Onaka, M. L. Patchett, J. Piel, M. J. Reaney, S. Rebuffat, R. P. Ross, H. G. Sahl, E. W. Schmidt, M. E. Selsted, K. Severinov, B. Shen, K. Sivonen, L. Smith, T. Stein, R. D. Sussmuth, J. R. Tagg, G. L. Tang, A. W. Truman, J. C. Vederas, C. T. Walsh, J. D. Walton, S. C. Wenzel, J. M. Willey and W. A. van der Donk (2013). “Ribosomally synthesized and post-translationally modified peptide natural products: overview and recommendations for a universal nomenclature.” Nat. Prod. Rep. 30(1): 108–160.

Bierbaum, G., H. Brotz, K. P. Koller and H. G. Sahl (1995). “Cloning, sequencing and production of the lantibiotic mersacidin.” FEMS Microbiol Lett 127(1-2): 121–126.

Bock, C., T. Zollmann, K.-A. Lindt, R. Tampé and R. Abele (2019). “Peptide translocation by the lysosomal ABC transporter TAPL is regulated by coupling efficiency and activation energy.” Scientific Reports 9(1): 11884.

Choudhury, H. G., Z. Tong, I. Mathavan, Y. Li, S. Iwata, S. Zirah, S. Rebuffat, H. W. van Veen and K. Beis (2014). “Structure of an antibacterial peptide ATP-binding cassette transporter in a novel outward occluded state.” Proc Natl Acad Sci U S A 111(25): 9145–9150.

Dischinger, J., S. Basi Chipalu and G. Bierbaum (2014). “Lantibiotics: promising candidates for future applications in health care.” Int J Med Microbiol 304(1): 51–62.

Dyballa, N. and S. Metzger (2009). “Fast and sensitive colloidal coomassie G-250 staining for proteins in polyacrylamide gels.” J Vis Exp(30).

Fath, M. J. and R. Kolter (1993). “ABC transporters: bacterial exporters.” Microbiol. Rev. 57(4): 995–1017.

Garg, N., L. M. Salazar-Ocampo and W. A. van der Donk (2013). “In vitro activity of the nisin dehydratase NisB.” Proc Natl Acad Sci U S A 110(18): 7258–7263.

Gross, E. and J. L. Morell (1967). “The presence of dehydroalanine in the antibiotic nisin and its relationship to activity.” J Am Chem Soc 89(11): 2791–2792.

Gross, E. and J. L. Morell (1968). “The number and nature of alpha, beta-unsaturated amino acids in nisin.” FEBS Lett 2(1): 61–64.

Havarstein, L. S., D. B. Diep and I. F. Nes (1995). “A family of bacteriocin ABC transporters carry out proteolytic processing of their substrates concomitant with export.” Mol Microbiol 16(2): 229–240.

Higgins, C. F. (1992). “ABC transporters: from microorganisms to man.” Annu Rev Cell Biol 8: 67–113.

Holo, H. and I. F. Nes (1989). “High-Frequency Transformation, by Electroporation, of *Lactococcus lactis subsp.* cremoris Grown with Glycine in Osmotically Stabilized Media.” Appl. Environ. Microbiol. 55(12): 3119–3123.

Hudson, G. A. and D. A. Mitchell (2018). “RiPP antibiotics: biosynthesis and engineering potential.” Curr Opin Microbiol 45: 61–69.

Infed, N., N. Hanekop, A. J. Driessen, S. H. Smits and L. Schmitt (2011). “Influence of detergents on the activity of the ABC transporter LmrA.” Biochim Biophys Acta 1808(9): 2313–2321.

Ingram, L. (1970). “A ribosomal mechanism for synthesis of peptides related to nisin.” Biochim Biophys Acta 224(1): 263–265.

Izaguirre, G. and J. N. Hansen (1997). “Use of alkaline phosphatase as a reporter polypeptide to study the role of the subtilin leader segment and the SpaT transporter in the posttranslational modifications and secretion of subtilin in Bacillus subtilis 168.” Appl Environ Microbiol 63(10): 3965–3971.

Jensen, P. R. and K. Hammer (1993). “Minimal Requirements for Exponential Growth of *Lactococcus lactis*.” Appl. Environ. Microbiol. 59(12): 4363–4366.

Karakas Sen, A., A. Narbad, N. Horn, H. M. Dodd, A. J. Parr, I. Colquhoun and M. J. Gasson (1999). “Post-translational modification of nisin. The involvement of NisB in the dehydration process.” Eur. J. Biochem. 261(2): 524–532.

Khusainov, R., R. Heils, J. Lubelski, G. N. Moll and O. P. Kuipers (2011). “Determining sites of interaction between prenisin and its modification enzymes NisB and NisC.” Mol Microbiol 82(3): 706–718.

Khusainov, R. and O. P. Kuipers (2013). “The presence of modifiable residues in the core peptide part of precursor nisin is not crucial for precursor nisin interactions with NisB- and NisC.” PLoS One 8(9): e74890.

Khusainov, R., G. N. Moll and O. P. Kuipers (2013). “Identification of distinct nisin leader peptide regions that determine interactions with the modification enzymes NisB and NisC.” FEBS Open Bio 3: 237–242.

Kiesau, P., U. Eikmanns, Z. Gutowski-Eckel, S. Weber, M. Hammelmann and K. D. Entian (1997). “Evidence for a multimeric subtilin synthetase complex.” J Bacteriol 179(5): 1475–1481.

Klein, C., C. Kaletta, N. Schnell and K. D. Entian (1992). “Analysis of genes involved in biosynthesis of the lantibiotic subtilin.” Appl Environ Microbiol 58(1): 132–142.

Kluskens, L. D., A. Kuipers, R. Rink, E. de Boef, S. Fekken, A. J. Driessen, O. P. Kuipers and G. N. Moll (2005). “Post-translational modification of therapeutic peptides by NisB, the dehydratase of the lantibiotic nisin.” Biochemistry 44(38): 12827–12834.

Koponen, O., M. Tolonen, M. Qiao, G. Wahlstrom, J. Helin and P. E. Saris (2002). “NisB is required for the dehydration and NisC for the lanthionine formation in the post-translational modification of nisin.” Microbiology 148(Pt 11): 3561–3568.

Kuipers, A., E. de Boef, R. Rink, S. Fekken, L. D. Kluskens, A. J. Driessen, K. Leenhouts, O. P. Kuipers and G. N. Moll (2004). “NisT, the transporter of the lantibiotic nisin, can transport fully modified, dehydrated, and unmodified prenisin and fusions of the leader peptide with non-lantibiotic peptides.” J. Biol. Chem. 279(21): 22176–22182.

Kuipers, A., J. Meijer-Wierenga, R. Rink, L. D. Kluskens and G. N. Moll (2008). “Mechanistic dissection of the enzyme complexes involved in biosynthesis of lacticin 3147 and nisin.” Appl Environ Microbiol 74(21): 6591–6597.

Kuipers, O. P., M. M. Beerthuyzen, R. J. Siezen and W. M. De Vos (1993). “Characterization of the nisin gene cluster nisABTCIPR of *Lactococcus lactis*. Requirement of expression of the nisA and nisI genes for development of immunity.” Eur. J. Biochem. 216(1): 281–291.

Laemmli, U. K. (1970). “Cleavage of structural proteins during the assembly of the head of bacteriophage T4.” Nature 227(5259): 680–685.

Lagedroste, M., J. Reiners, S. H. J. Smits and L. Schmitt (2019). “Systematic characterization of position one variants within the lantibiotic nisin.” Sci Rep 9(1): 935.

Lenders, M. H., T. Beer, S. H. Smits and L. Schmitt (2016). “In vivo quantification of the secretion rates of the hemolysin A Type I secretion system.” Sci Rep 6: 33275.

Li, B. and W. A. van der Donk (2007). “Identification of essential catalytic residues of the cyclase NisC involved in the biosynthesis of nisin.” J Biol Chem 282(29): 21169–21175.

Li, B., J. P. Yu, J. S. Brunzelle, G. N. Moll, W. A. van der Donk and S. K. Nair (2006). “Structure and mechanism of the lantibiotic cyclase involved in nisin biosynthesis.” Science 311(5766): 1464–1467.

Lin, D. Y., S. Huang and J. Chen (2015). “Crystal structures of a polypeptide processing and secretion transporter.” Nature 523(7561): 425–430.

Lubelski, J., R. Khusainov and O. P. Kuipers (2009). “Directionality and coordination of dehydration and ring formation during biosynthesis of the lantibiotic nisin.” J Biol Chem 284(38): 25962–25972.

Mavaro, A., A. Abts, P. J. Bakkes, G. N. Moll, A. J. Driessen, S. H. Smits and L. Schmitt (2011). “Substrate recognition and specificity of the NisB protein, the lantibiotic dehydratase involved in nisin biosynthesis.” J Biol Chem 286(35): 30552–30560.

Meyer, C., G. Bierbaum, C. Heidrich, M. Reis, J. Suling, M. I. Iglesias-Wind, C. Kempter, E. Molitor and H. G. Sahl (1995). “Nucleotide sequence of the lantibiotic Pep5 biosynthetic gene cluster and functional analysis of PepP and PepC. Evidence for a role of PepC in thioether formation.” Eur. J. Biochem. 232(2): 478–489.

Morgan, J. L. W., J. F. Acheson and J. Zimmer (2017). “Structure of a Type-1 Secretion System ABC Transporter.” Structure 25(3): 522–529.

Nagao, J., Y. Aso, T. Sashihara, K. Shioya, A. Adachi, J. Nakayama and K. Sonomoto (2005). “Localization and interaction of the biosynthetic proteins for the lantibiotic, Nukacin ISK-1.” Biosci Biotechnol Biochem 69(7): 1341–1347.

Newman, D. J. and G. M. Cragg (2016). “Natural Products as Sources of New Drugs from 1981 to 2014.” J Nat Prod 79(3): 629–661.

Newton, G. G., E. P. Abraham and N. J. Berridge (1953). “Sulphur-containing amino-acids of nisin.” Nature 171(4353): 606.

Okeley, N. M., M. Paul, J. P. Stasser, N. Blackburn and W. A. van der Donk (2003). “SpaC and NisC, the cyclases involved in subtilin and nisin biosynthesis, are zinc proteins.” Biochemistry 42(46): 13613–13624.

Ortega, M. A., Y. Hao, Q. Zhang, M. C. Walker, W. A. van der Donk and S. K. Nair (2015). “Structure and mechanism of the tRNA-dependent lantibiotic dehydratase NisB.” Nature 517(7535): 509–512.

Quiao, M. and P. E. Saris (1996). “Evidence for a role of NisT in transport of the lantibiotic nisin produced by Lactococcus lactis N8.” FEMS Microbiol Lett 144(1): 89–93.

Ra, S. R., M. Qiao, T. Immonen, I. Pujana and E. J. Saris (1996). “Genes responsible for nisin synthesis, regulation and immunity form a regulon of two operons and are induced by nisin in Lactoccocus lactis N8.” Microbiology 142 (Pt 5): 1281–1288.

Reimann, S., G. Poschmann, K. Kanonenberg, K. Stuhler, S. H. Smits and L. Schmitt (2016). “Interdomain regulation of the ATPase activity of the ABC transporter haemolysin B from Escherichia coli.” Biochem J 473(16): 2471–2483.

Reiners, J., A. Abts, R. Clemens, S. H. Smits and L. Schmitt (2017). “Stoichiometry and structure of a lantibiotic maturation complex.” Sci Rep 7: 42163.

Repka, L. M., K. J. Hetrick, S. H. Chee and W. A. van der Donk (2018). “Characterization of Leader Peptide Binding During Catalysis by the Nisin Dehydratase NisB.” J Am Chem Soc 140(12): 4200–4203.

Rink, R., L. D. Kluskens, A. Kuipers, A. J. Driessen, O. P. Kuipers and G. N. Moll (2007). “NisC, the cyclase of the lantibiotic nisin, can catalyze cyclization of designed nonlantibiotic peptides.” Biochemistry 46(45): 13179–13189.

Rink, R., A. Kuipers, E. de Boef, K. J. Leenhouts, A. J. Driessen, G. N. Moll and O. P. Kuipers (2005). “Lantibiotic structures as guidelines for the design of peptides that can be modified by lantibiotic enzymes.” Biochemistry 44(24): 8873–8882.

Robson, A., V. A. M. Gold, S. Hodson, A. R. Clarke and I. Collinson (2009). “Energy transduction in protein transport and the ATP hydrolytic cycle of SecA.” Proceedings of the National Academy of Sciences 106(13): 5111–5116.

Schagger, H. (2006). “Tricine-SDS-PAGE.” Nat. Protoc. 1(1): 16–22.

Schneider, C. A., W. S. Rasband and K. W. Eliceiri (2012). “NIH Image to ImageJ: 25 years of image analysis.” Nat Methods 9(7): 671–675.

Schnell, N., G. Engelke, J. Augustin, R. Rosenstein, V. Ungermann, F. Gotz and K. D. Entian (1992). “Analysis of genes involved in the biosynthesis of lantibiotic epidermin.” Eur J Biochem 204(1): 57–68.

Siegers, K., S. Heinzmann and K. D. Entian (1996). “Biosynthesis of lantibiotic nisin. Posttranslational modification of its prepeptide occurs at a multimeric membrane-associated lanthionine synthetase complex.” J. Biol. Chem. 271(21): 12294–12301.

Terzaghi, B. E. and W. E. Sandine (1975). “Improved medium for lactic streptococci and their bacteriophages.” Appl. Microbiol. 29(6): 807–813.

van den Berg van Saparoea, H. B., P. J. Bakkes, G. N. Moll and A. J. Driessen (2008). “Distinct contributions of the nisin biosynthesis enzymes NisB and NisC and transporter NisT to prenisin production by Lactococcus lactis.” Appl Environ Microbiol 74(17): 5541–5548.

van der Meer, J. R., J. Polman, M. M. Beerthuyzen, R. J. Siezen, O. P. Kuipers and W. M. De Vos (1993). “Characterization of the *Lactococcus lactis* nisin A operon genes nisP, encoding a subtilisin-like serine protease involved in precursor processing, and nisR, encoding a regulatory protein involved in nisin biosynthesis.” J. Bacteriol. 175(9): 2578–2588.

van der Meer, J. R., H. S. Rollema, R. J. Siezen, M. M. Beerthuyzen, O. P. Kuipers and W. M. de Vos (1994). “Influence of amino acid substitutions in the nisin leader peptide on biosynthesis and secretion of nisin by *Lactococcus lactis*.” J. Biol. Chem. 269(5): 3555–3562.

van Heel, A. J., T. G. Kloosterman, M. Montalban-Lopez, J. Deng, A. Plat, B. Baudu, D. Hendriks, G. N. Moll and O. P. Kuipers (2016). “Discovery, Production and Modification of Five Novel Lantibiotics Using the Promiscuous Nisin Modification Machinery.” ACS Synth Biol 5(10): 1146–1154.

Zaitseva, J., S. Jenewein, T. Jumpertz, I. B. Holland and L. Schmitt (2005). “H662 is the linchpin of ATP hydrolysis in the nucleotide-binding domain of the ABC transporter HlyB.” EMBO J 24(11): 1901–1910.

Zheng, S., J. I. Nagao, M. Nishie, T. Zendo and K. Sonomoto (2017). “ATPase activity regulation by leader peptide processing of ABC transporter maturation and secretion protein, NukT, for lantibiotic nukacin ISK-1.” Appl Microbiol Biotechnol.

Zhou, L., A. J. van Heel and O. P. Kuipers (2015). “The length of a lantibiotic hinge region has profound influence on antimicrobial activity and host specificity.” Front Microbiol 6: 11.

Zhou, L., A. J. van Heel, M. Montalban-Lopez and O. P. Kuipers (2016). “Potentiating the Activity of Nisin against Escherichia coli.” Front Cell Dev Biol 4: 7.

